# Ontology-aware deep learning for antibiotic resistance gene prediction: novel function discovery and comprehensive profiling from metagenomic data

**DOI:** 10.1101/2021.07.30.454403

**Authors:** Yuguo Zha, Cheng Chen, Qihong Jiao, Xiaomei Zeng, Xuefeng Cui, Kang Ning

## Abstract

Antibiotic resistance genes (ARGs) have emerged in pathogens and arousing a worldwide concern, which is estimated to cause millions of deaths each year globally. Accurately identifying and classifying ARGs is a formidable challenge in studying the generation and spread of antibiotic resistance. Current methods could identify close homologous ARGs, have limited utility for discovery of novel ARGs, thus rendering the profiling of ARGs incomprehensive. Here, an ontology-aware neural network (ONN) approach, ONN4ARG, is proposed for comprehensive ARG discovery. Systematic evaluation shows ONN4ARG is advanced than previous methods such as DeepARG in efficiency, accuracy, and comprehensiveness. Experiments using 200 million candidate microbial genes collected from 815 microbial community samples from diverse environments or hosts have resulted in 120,726 candidate ARGs, out of which more than 20% are not yet present in public databases. These comprehensive set of ARGs have clarified the environment-specific and host-specific patterns. The wet-experimental functional validation, together with structural investigation of docking sites, have also validated a novel streptomycin resistance gene from oral microbiome samples, confirming ONN4ARG’s ability for novel ARGs identification. In summary, ONN4ARG is superior to existing methods in efficiency, accuracy, and comprehensiveness. It enables comprehensive ARG discovery, which is helpful towards a grand view of ARGs worldwide. ONN4ARG is available at https://github.com/HUST-NingKang-Lab/ONN4ARG, and online web service is available at http://onn4arg.xfcui.com/.

## Introduction

With the development of metagenomics and next-generation sequencing, many new microbial taxa and genes have been discovered, but different kinds of “unknowns” remain. For instance, the microbes found in the human gut microbiome involve 25 phyla, more than 2,000 genera, and 5,000 species [1]. However, the functional diversity of microbiomes has not been fully explored, and about 40% of microbial gene functions remain to be discovered [2]. A typical example is the antibiotic resistance gene (ARG), which is an urgent and growing threat to public health [3]. In the past few decades, problems caused by antibiotic resistance have drawn the public’s attention [4]. Antimicrobial resistance genomic data is an ever-expanding data source, with many new ARG families discovered in recent years [5, 6]. The discovery of resistance genes in diverse environments offers possibilities for early surveillance, actions to reduce transmission, gene-based diagnostics, and improved treatment [7].

Existing annotated ARGs have been curated manually or automatically for decades. Presently, there are 4,661 annotated ARGs in the reference database CARD [5, 6] (v3.2.5, released in September 2022), 3,131 in the ResFinder database [8] (as of December 2022), and 2,476 in SwissProt [9] (as of December 2022). These annotated ARGs are categorized into antibiotic resistance types, which are organized in an ontology structure (**Methods**, **Supplementary Figure S1**), in which higher-level ARG types cover lower-level ARG types. Current ARG databases are far from complete: though no ARG database contains more than 4,000 well-annotated ARGs, NCBI non-redundant database searches yielded more than 7,000 putative genes annotated with “antibiotic resistance” as of May 2021. Therefore, we deemed that there is a large gap between the genes annotated in ARG databases and the possible ARGs that already exist in general databases, not to mention ARGs that are not yet annotated.

Many ARG prediction tools have been proposed in the past few years [8, 10–20]. These tools can generally be divided into two approaches. One approach is sequence-alignment, such as BLAST [21], USEARCH [22], and Diamond [23], which uses homologous genes to annotate unclassified genes. A confident prediction requires a homolog with sequence identity greater than 80% in many programs, such as ResFinder [8, 11]. The other approach is deep learning, such as DeepARG [12] and HMD-ARG [16], which uses neural network models to predict and annotate ARGs.

Several limitations still preclude comprehensive profiling of ARGs. A more comprehensive set of ARGs could be roughly defined as having more ARGs in type and number with less false-positive entries, regardless of the homology with known ARGs, and many of these ARGs could be experimentally validated. Based on this definition, existing tools fall short in comprehensive profiling of ARGs. First, existing tools are limited to a few types of ARGs due to the fact that the datasets used for building models are specialized. For example, HMD-ARG [16] identifies only 15 types of resistance genes, and PATRIC [13] is limited to identifying ARGs encoding resistance to carbapenem, methicillin, and beta-lactam antibiotics. Second, existing tools fall short in discovering novel ARGs, which usually lack homology to known sequences in the reference databases. For instance, the gene POCOZ1 (VraR) that confers resistance to vancomycin has a sequence identity of only 24% to the homolog from the CARD [12]. Therefore, there is an urgent need for a new approach to address these limitations.

Here, we propose an ontology-aware deep learning approach, ONN4ARG, which allows comprehensive identification of ARGs. Systematic evaluation based on the ONN4ARG-DB, CARD, and ResFinder datasets shows that the ONN4ARG model outperforms state-of-the-art models such as DeepARG, especially for the detection of remotely homologous ARGs. Experiments based on more than 200 million candidate microbial genes collected from 815 samples in various environments have resulted in 120,726 candidate ARGs, out of which more than 20% are not yet present in public databases. Our experiments confirmed that ARGs are both environment-specific and host-specific, exemplified by the rifamycin resistance genes which are enriched in Actinobacteria and in soil environment. Case study of a recently experimentally validated ARG gene GAR [7] have also verified the ability of ONN4ARG for novel ARG discovery. We also validated a novel streptomycin resistance gene from oral microbiome samples by wet-lab experiment. In summary, ONN4ARG enables comprehensive ARG discovery, which provides a relatively complete picture of the prevalence of ARGs, as well as leads a way towards a grand view of ARGs worldwide.

## Results

### ONN4ARG model employs an ontology-aware neural network for ARG identification and classification

To address the large gap between the genes annotated in ARG databases and the possible ARGs that already exist in general databases along with the ARGs that are not yet annotated, we propose ONN4ARG, which is an ontology-aware neural network model (**Figure 1**, **Supplementary Figure S1**) that predict ARGs in a comprehensive manner. ONN4ARG takes similarities (e.g., identity, e-value, bit-score) between the query gene sequence and ARG gene sequences and profiles (i.e., PSSM) as inputs and predicts ARG annotations (**Figure 1B**). These sequence-alignment similarities and profile-alignment similarities are pre-processed by calling Diamond [23] and HHblits [24]. ONN4ARG generates hierarchical annotations of antibiotic resistance types, which are compatible with the antibiotic resistance ontology structure (**Figure 1A, C**). One advantage of ONN4ARG over state-of-the-art models is that ONN4ARG employs a novel ontology-aware layer that incorporates ancestor and descendent annotations to enhance annotation accuracies (**Methods**). To train and evaluate our ONN4ARG model and for rapid deployment of ARG discovery in multiple contexts, we also built an ARG database (**Figure 1D**), namely, ONN4ARG-DB, which comprises ARGs from CARD and UniProt (see **Methods**).

**Figure 1.**
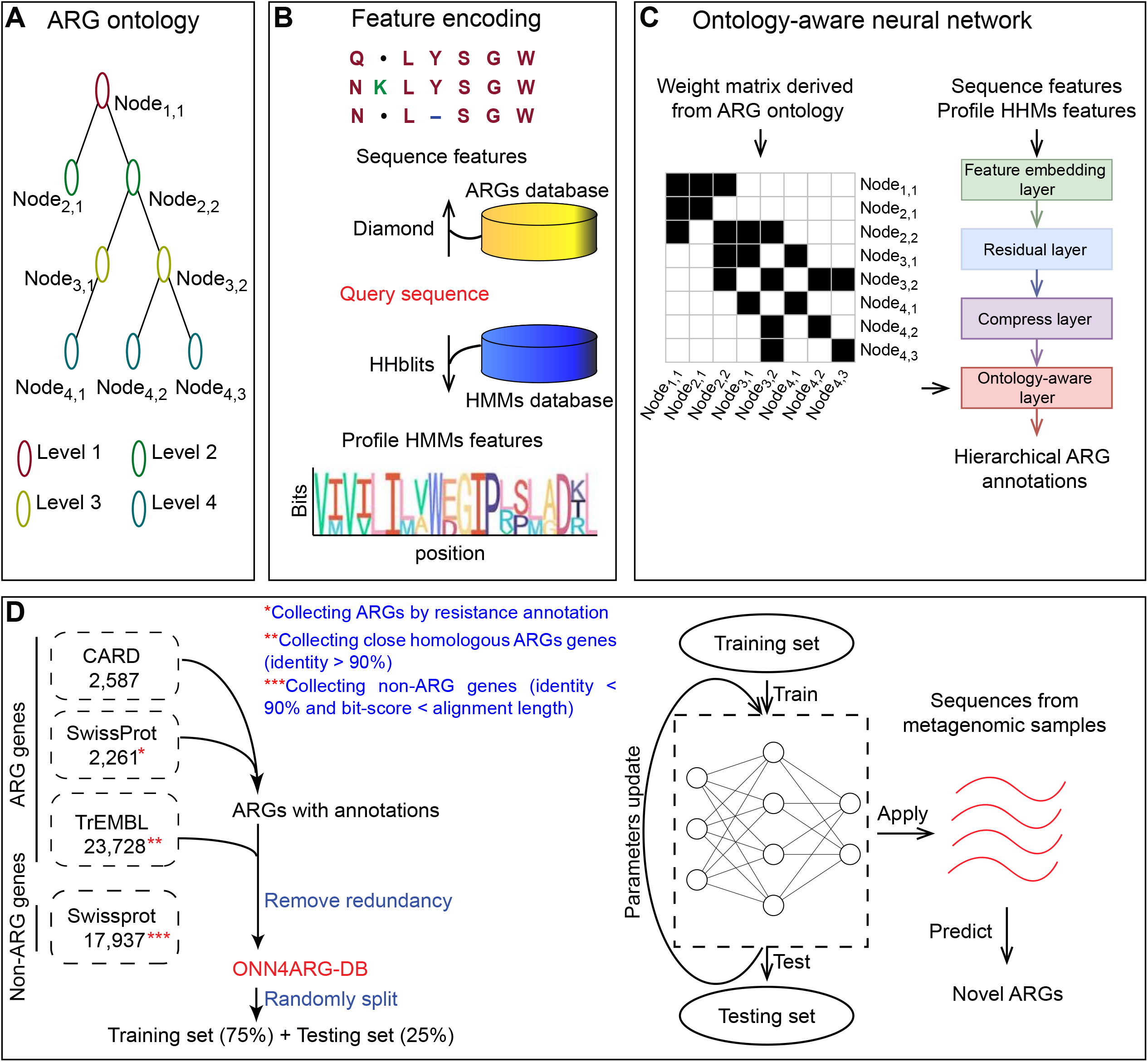
Overview of the ONN4ARG model and its use for novel ARG discovery. (**A**) The antibiotic resistance gene ontology contains four levels. The root (first level) is a single node, namely, “arg”. There are 1, 2, 34, and 277 nodes from the first level to the fourth level, respectively. (**B**) The feature encoding procedure of ONN4ARG model. The sequence alignment features and profile HMMs features are encoded by calling Diamond and HHblits. (**C**) The architecture of the ontology-aware neural network could be described in four functional layers, including feature embedding layer, residual layer, compress layer and ontology-aware layer. The ontology-aware layer is a partially connected layer which encourage annotation predictions satisfying the ontology rules (i.e., the ontology tree structure). Specially, weight between nodes with relationship (e.g., parent and child) satisfying the ontology rules would be saved in the partially connected layer, and weights between irrelevant nodes would be masked. (**D**) Building the dataset for training and testing, and applying ONN4ARG model on metagenomic samples to discover candidate novel ARGs.

### Systematic evaluation and comparison

Systematic evaluation based on the ONN4ARG-DB showed our model’s high efficiency, high accuracy, and comprehensiveness for ARG identification. ONN4ARG is fast since it could complete ARG identification for all genes in the testing dataset within four hours, which is equivalent to one second per gene identification. As shown in **Figure 2A**, ONN4ARG was more accurate for ARG identification (overall accuracy of 97.70%, **Table 1**) compared to sequence alignment (overall accuracy of 69.11%), and ONN4ARG has a slight advantage over DeepARG (overall accuracy of 96.39%). Moreover, ONN4ARG achieved an overall precision of 75.59% and an overall recall of 89.93%, which were higher than DeepARG’s overall precision of 68.30% and overall recall of 77.84% (**Figure 2B**, **Table 2**). It is natural that ONN4ARG could not outperform DeepARG in all resistance types and this is exemplified by results on pleuromutilin due to the small number of sequences for pleuromutilin in the ONN4ARG-DB. ONN4ARG demonstrates an advantage over other methods in identification of remotely homologous ARGs whose sequences are not similar to existing ARG sequences (**Tables 2** and **3**). In this context, when testing with only remotely homologs (i.e., the masking threshold of testing set equal to 0.4, see **Methods**), ONN4ARG achieves an accuracy of 94.26%, which is largely improved from 89.85% of DeepARG. These results validate ONN4ARG’s better generalization abilities than sequence-alignment and DeepARG, which makes ONN4ARG especially suitable for identification of remotely homologous ARGs and indicates ONN4ARG’s ability for novel ARG discovery (**Tables 1–3**).

**Figure 2.**
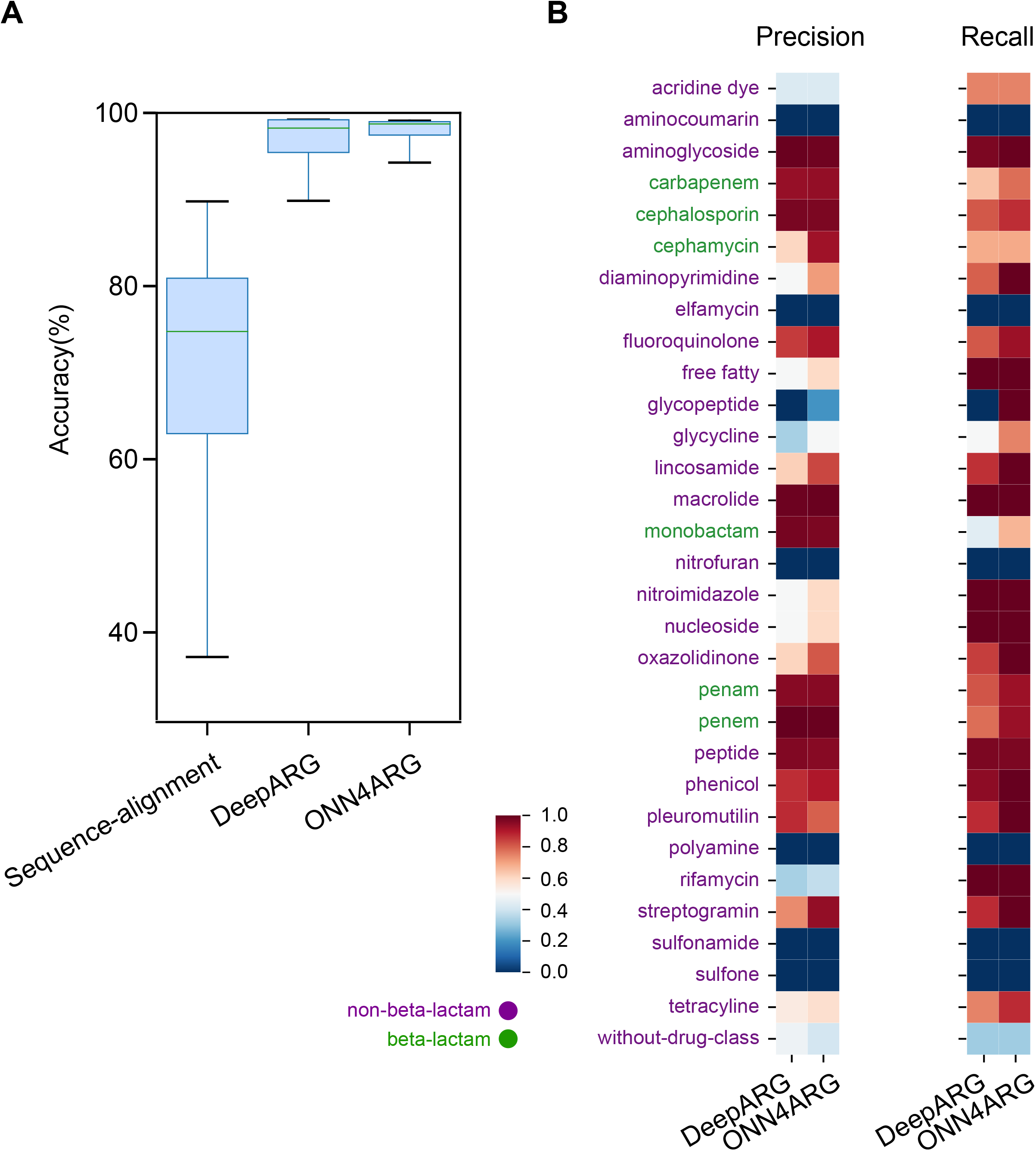
Systematic evaluation and comparison between sequence-alignment, DeepARG, and ONN4ARG. (**A**) The accuracy of three models on ARG classification was assessed using a box plot. Diamond was used for sequence-alignment; significance test was based on the *t*-test. (**B**) The precision and recall of DeepARG and ONN4ARG on ARG classification for each antibiotic resistance type. The masking threshold of testing set equaled 0.4 (details of masking threshold are provided in **Methods**).

We have also tested ONN4ARG on a verification set built from the CARD database version 3.1.3. Results showed our model outperformed other methods in terms of accuracy and efficiency, i.e., high accuracy and less time usage, given that the memory usage is acceptable for a regular laptop (**Supplementary Table S1**). We have also evaluated ONN4ARG on the ResFinder database version 4.1, which involves thousands of manually curated ARGs [8]. Results showed that ONN4ARG achieved an accuracy higher than 90% for most types of resistance, while DeepARG was less accurate than ONN4ARG, except for the fosfomycin resistance (**Supplementary Table S2**).

### Applications of ONN4ARG on metagenomic data

We collected metagenomic samples from several published studies [25, 26]. These samples were mainly from “marine,” “soil,” and “human” environments. Human-associated samples consisted of two gut groups (one group from Madagascar, i.e., GutM; the other group from Denmark, i.e., GutD), one oral group, and one skin group (both oral and skin groups were from the HMP project). For details on these samples, see **Supplementary Table S3**. Then, genes were obtained by calling Prodigal [27] with default parameters. The ONN4ARG model was used to predict whether these unclassified genes were ARGs and their corresponding resistance types. In total, 120,726 ARGs were identified from microbiome samples, many of which are novel, which greatly expands the existing ARG repositories.

### Broad-spectrum profile of predicted ARGs among diverse environments

We investigated the broad-spectrum profile of these predicted ARGs among diverse environments. First, we investigated the proportion of predicted ARGs for different sequence lengths. The distribution shows that about half of the predicted ARGs have a length of 128–256 amino acid residues (**Figure 3A**). We also analyzed the protein domain of these predicted ARGs by searching the conserved domain database (CDD, last update Aug 2022) using RPS-BLAST tool version 2.9.0. Results showed that most of these predicted ARGs (over 97%) have protein domains that resemble those with known catalytic activity and/or may bind to the antimicrobials they are predicted to elicit resistance against (**Supplementary Table S4**). Second, we found that human-associated microbiome samples carry a higher abundance of ARGs, especially for the oral group, in which more than one resistance gene could be observed out of a hundred genes on average (**Figure 3B**, **Supplementary Table S5**). Third, we tested the novelty of these predicted ARGs. We found that about a third of them (42,848 out of all 120,726 ARGs) had sequence identity of less than 40% to their homologs in the ONN4ARG-DB (**Figure 3C**). We define these ARGs as candidate novel ARGs, which have low sequence identities when aligned to their homologs in the reference database (i.e., ONN4ARG-DB). For example, we found 45% of predicted ARGs in the marine group were candidate novel ARGs (**Figure 3C**).

**Figure 3.**
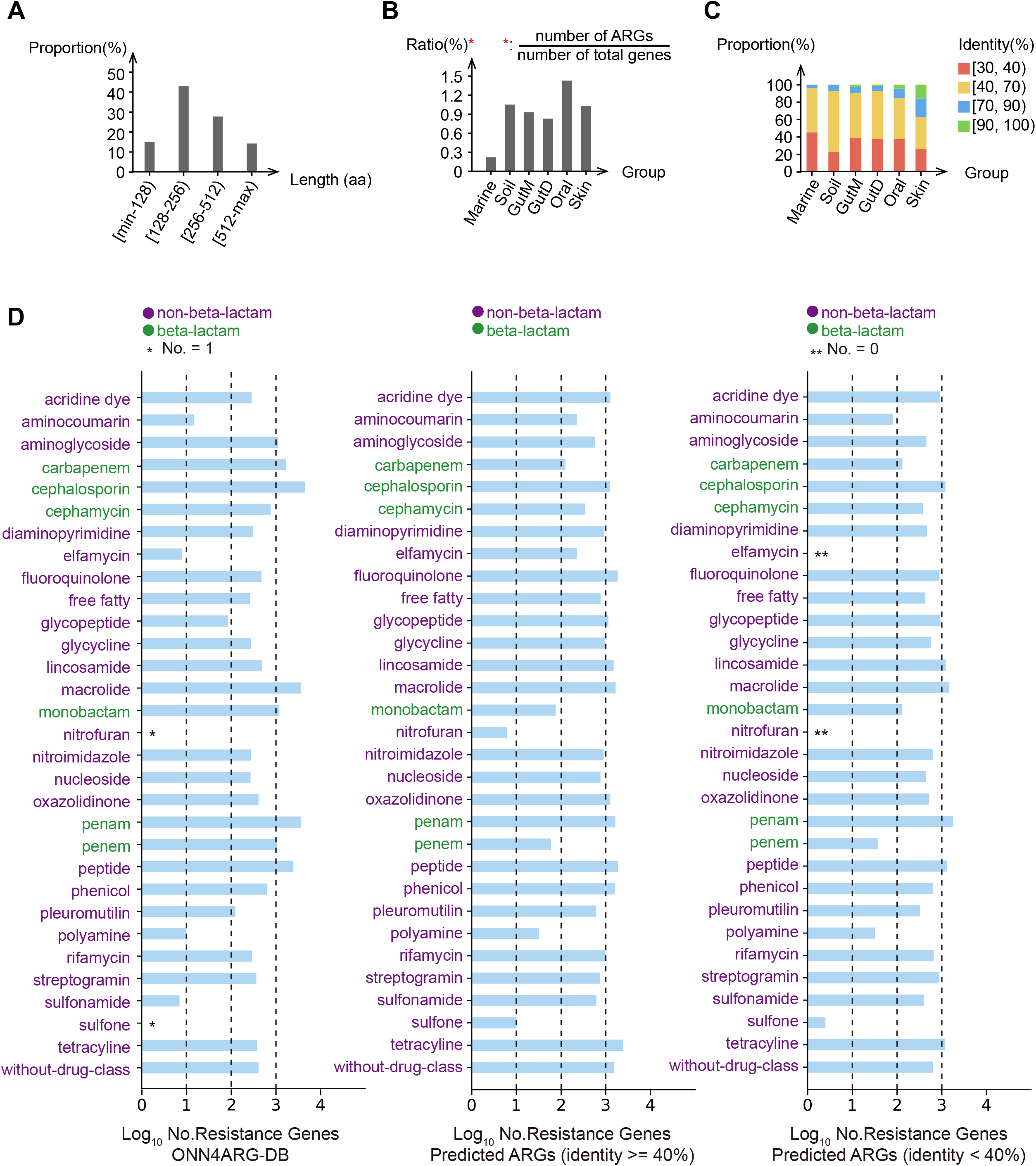
Broad-spectrum profile of predicted ARGs among diverse environments. (**A**) The proportion of predicted ARGs for different protein sequence lengths. (**B**) The abundance ratio of predicted ARGs among diverse environments. Abundance ratio was defined as the number of ARGs divided by the number of total genes. (**C**) The proportion of predicted ARGs for different sequence identities among diverse environments. (**D**) Number of genes in ONN4ARG-DB (left), predicted homologous ARGs (middle), and predicted novel ARGs (right) for various resistance types. The horizontal axis indicates the logarithmic number of genes, and the vertical axis indicates different antibiotic resistance types. We collected metagenomic samples from several published studies; these samples were mainly from “marine,” “soil,” and “human” environments. Human-associated samples consisted of two gut groups (one group from Madagascar, i.e., GutM; the other group from Denmark, i.e., GutD), one oral group, and one skin group (both oral and skin groups were from the HMP project).

In total, 31 ARG types were detected in these various environments (**Figure 3D**, **Supplementary Figure S2**). The number of predicted ARG sequences for different types varied greatly, from a few (i.e., nitrofuran) to thousands (i.e., fluoroquinolone). In general, fluoroquinolone and tetracycline resistance genes were more abundant than other types (**Figure 3D**). As expected, these abundant ARGs were usually associated with the antibiotics used extensively in human medicine or veterinary medicine, including growth promotion [28].

### Enrichment of predicted ARGs among diverse hosts and environments

Rapid deciphering of potential antimicrobial-resistant pathogens is necessary for effective public health monitoring. The host-tracking of ARGs allows for accurate identification of pathogens. Therefore, we conducted taxonomy analysis to track the hosts of these predicted ARGs by using Kraken2 [29]. Results showed that there are 949 genera, each genus carries at least one type of ARG (**Supplementary Table S6**). The host composition and distribution of all classified ARGs for the most abundant 20 genera are displayed in **Supplementary Figure S3**. The host distribution shows that these ARGs are primarily affiliated with Proteobacteria (38.2%). The most abundant ARGs carried by the 20 genera were resistance types of fluoroquinolone, macrolide, peptide, penam, and tetracycline, accounting for about half of the total ARGs. Network inference based on strong (Spearman’s *ρ* > 0.8) and significant (Welch’s *t*-test, P-value < 0.01) correlations showed the co-occurrence patterns among ARGs and microbial taxa (**Supplementary Figure S4**, **Supplementary File S1**). For example, ARGs that belong to beta-lactam resistance type (e.g., cephamycin, penam, penem, and monobactam) were observed to appear together in Proteobacteria.

Enrichment analyses showed that ARGs are both environment-specific and host-specific (**Figure 4**). We found that the proportion of certain types of ARGs was higher in certain environments than in others. For example, rifamycin resistance genes were found enriched in the soil environment (with proportion of 0.1%) and enriched in the Actinobacteria (with proportion of 4.7%) (**Figure 4**). Rifamycin is an important antibacterial agent active against gram-positive bacteria, and it has a wide range of applications [30, 31]. The enrichment results were not surprising because *Actinomycetes* is a representative genus widely distributed in various soil environments, and its rifamycin resistance is compatible with its ability for rifamycin production [32–35].

**Figure 4.**
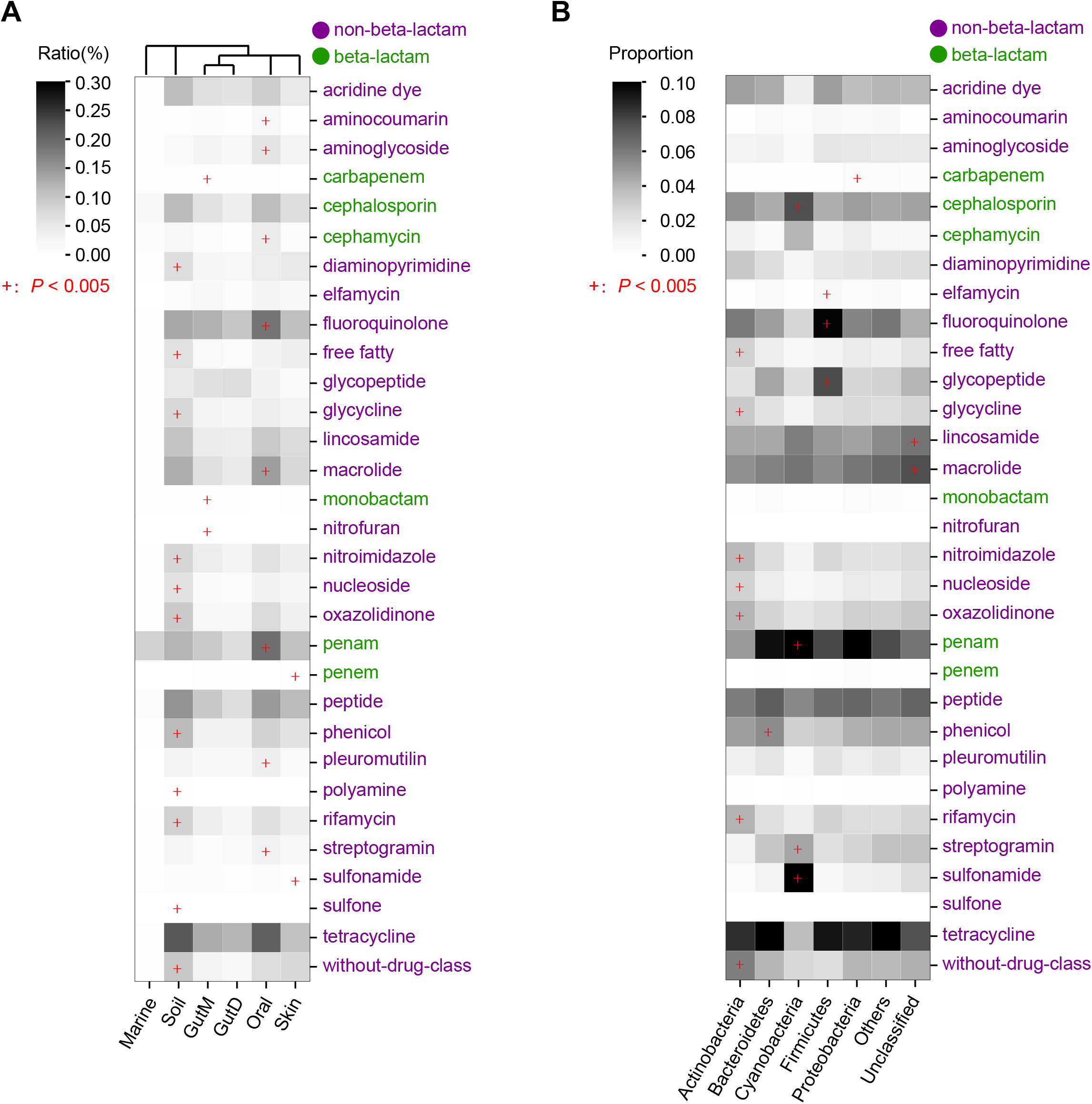
Enrichment of predicted ARGs among diverse environments and hosts. (**A**) Relative abundance and enrichment of ARGs among diverse environments. Abundance ratio was defined as the number of ARGs divided by the number of total genes. (**B**) Proportion and enrichment of ARGs among diverse hosts. Colors indicate the proportion of ARGs for each phylum and resistance type. Results for the most abundant five phyla that carry ARGs are shown. “+”: P-value < 0.005 (Welch’s *t*-test, one-tailed).

### Evaluation of the ability for novel ARG identification using a recently annotated ARG

We further evaluated ONN4ARG’s ability for novel ARG identification based on a newly annotated aminoglycoside resistance gene, GAR, which has been reported in a previous study by Böhm et al [7]. GAR is a recently reported aminoglycoside resistance gene, which is not present in CARD (v3.2.5), UniProt (as of December 2022), DEEPARG-DB (v1.0.2), HMD-ARG-DB (as of December 2022), and ONN4ARG-DB. We searched the sequence of GAR with both DeepARG and HMD-ARG models, and the results showed that both of these models indicated it as non-ARG. We searched the sequence of GAR against all the sequences in ONN4ARG-DB using Diamond and did not find any homologous gene as well. Reassuringly, the prediction by ONN4ARG identified GAR as an ARG resistant to non-beta-lactam with high confidence (probability score = 100%). We should emphasize that though ONN4ARG predict GAR as non-beta-lactam and not as sub-type of aminoglycoside, ONN4ARG can give information about ancestors (or categories at higher levels) of the novel ARG, provide clues about novel knowledge.

### Functional verification of candidate novel resistance genes

To identify promising putative novel resistance genes, we used four criteria: (i) remotely homologs to reference ARGs, (ii) prediction with high confidence, (iii) predicted to be single-type resistance, and (iv) the host is known. Despite the large number of candidate genes discovered by the ONN4ARG model, only 4,365 ARGs fulfilled all mentioned criteria (**Supplementary Table S7**).

To showcase the actual function of the predicted ARGs, we analyzed tens of ARGs belonging to the streptomycin resistance, and all of these ARGs have high confidence predicted by the ONN4ARG model. The experiment results showed that the Candi_60363_1 is one of the most promising ARG, which showed a high minimal inhibitory concentration (MIC) compared to negative control. Thus, we selected the Candi_60363_1 for further experimental validation (**Supplementary Table S8** and **S9**). Candi_60363_1, detected in *Streptococcus* in the oral environment, was predicted to confer resistance to streptomycin (belonging to aminoglycoside). One positive control from CARD (AHE40557.1, streptomycin resistance) was used for verification of the experimental system. All these genes were heterologously expressed in the *E. coli* BL21 (DE3) host by the induction of Isopropyl β-D-1-thiogalactopyranoside (IPTG) and tested for minimal inhibitory concentration (MIC) (**Figure 5A**). Results showed that the mRNA level of the genes increased with the addition of 1 mM IPTG compared with that without IPTG (**Figure 5B**), which verified the expression of the genes induced by IPTG. Furthermore, the MIC of the strain containing the positive control gene AHE40557.1 was more than 1,024 μg/ml (**Supplementary Figure S5**), which is consistent with previous reports [36, 37]. This verified that our MIC measuring experimental system works well. Our results showed that the MIC of the strain containing Candi_60363_1 was significantly higher than the negative control containing no insert (Welch’s t-test, one-tailed, P-value = 3.5e-3), which demonstrated the increased resistance to streptomycin of the novel candidate gene Candi_60363_1 (**Figure 5C**, **Supplementary Figure S5**).

**Figure 5.**
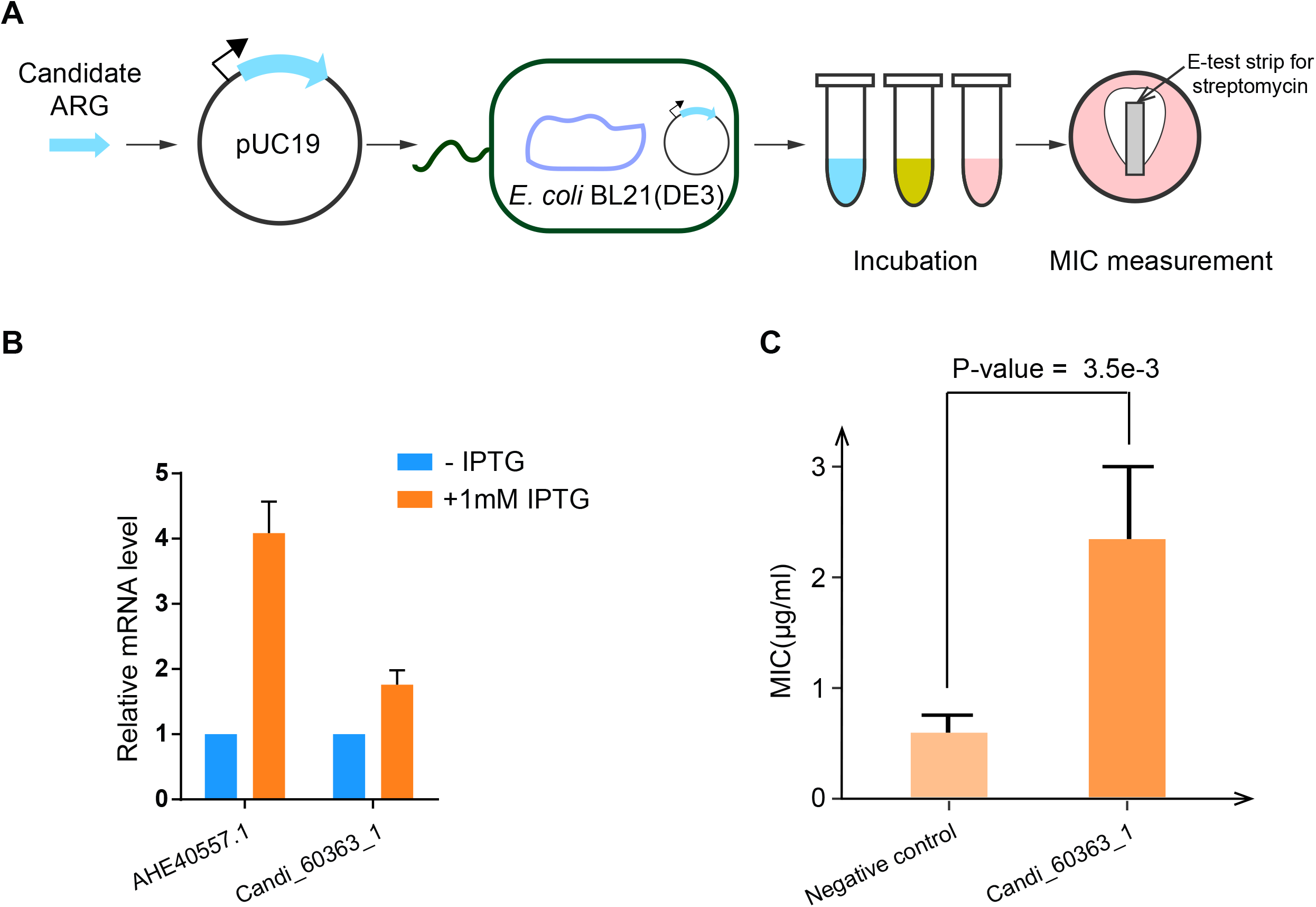
Functional validation of a predicted candidate novel ARG. (**A**) A diagram showing the procedure of heterologous expression and functional analysis of the predicted candidate ARG in the *E. coli* BL21 (DE3) host. (**B**) Gene expression validation of the predicted candidate ARG. The vertical axis indicates the relative mRNA level. (**C**) The MIC of the predicted candidate ARG and negative control. The vertical axis indicates the MIC value. The MIC of the predicted candidate novel ARG is significantly higher than the negative control (Welch’s *t*-test, one-tailed, P-value = 3.5e-3).

### Phylogeny and structure of Candi_60363_1

There are remotely similarities between Candi_60363_1 and all known ARGs in the reference database, including aminoglycoside resistance genes. The InterPro search results showed the protein family matching to Candi_60363_1 is IPR007530, which is also known as aminoglycoside 6-adenylyltransferase that confers resistance to aminoglycoside antibiotics. Then, we used BLAST to search homologs of Candi_60363_1 from the NCBI non-redundant protein database. The BLAST result showed that there are 44 homologs with sequence identity greater than 80%, and they are from various organisms (**Supplementary Table S10**), such as *Streptococcus oralis, Peptoniphilus lacrimalis DNF00528*, and *Mycobacteroides abscessus subsp. Abscessus*. Considering that Candi_60363_1 is harbored by distantly related species, it obviously has mobility. Notably, the most similar protein of Candi_60363_1 from the NCBI non-redundant protein database (87.5% identity, SHZ78752.1) is also annotated as aminoglycoside adenylyltransferase (**Supplementary Table S10**). Taken together, Candi_60363_1 is highly likely to be an ARG that confers resistance to aminoglycoside antibiotics.

Aminoglycoside modifying enzymes are the most clinically important resistance mechanism against aminoglycosides [38]. They are divided into three enzymatic classes, namely, aminoglycoside N-acetyltransferase (AAC), O-nucleotidyltransferase (ANT), and O-phosphotransferase (APH). We investigated the phylogenetic relationship between Candi_60363_1 and the known aminoglycoside modifying enzymes. The phylogenetic tree of Candi_60363_1 and related proteins (**Figure 6A**) shows that Candi_60363_1 is clearly separated from the known aminoglycoside modifying enzymes and is located among proteins mostly annotated as aminoglycoside adenylyltransferase. Phylogenetic analysis indicated its evolutionarily close relationships with known aminoglycoside adenylyltransferase.

**Figure 6.**
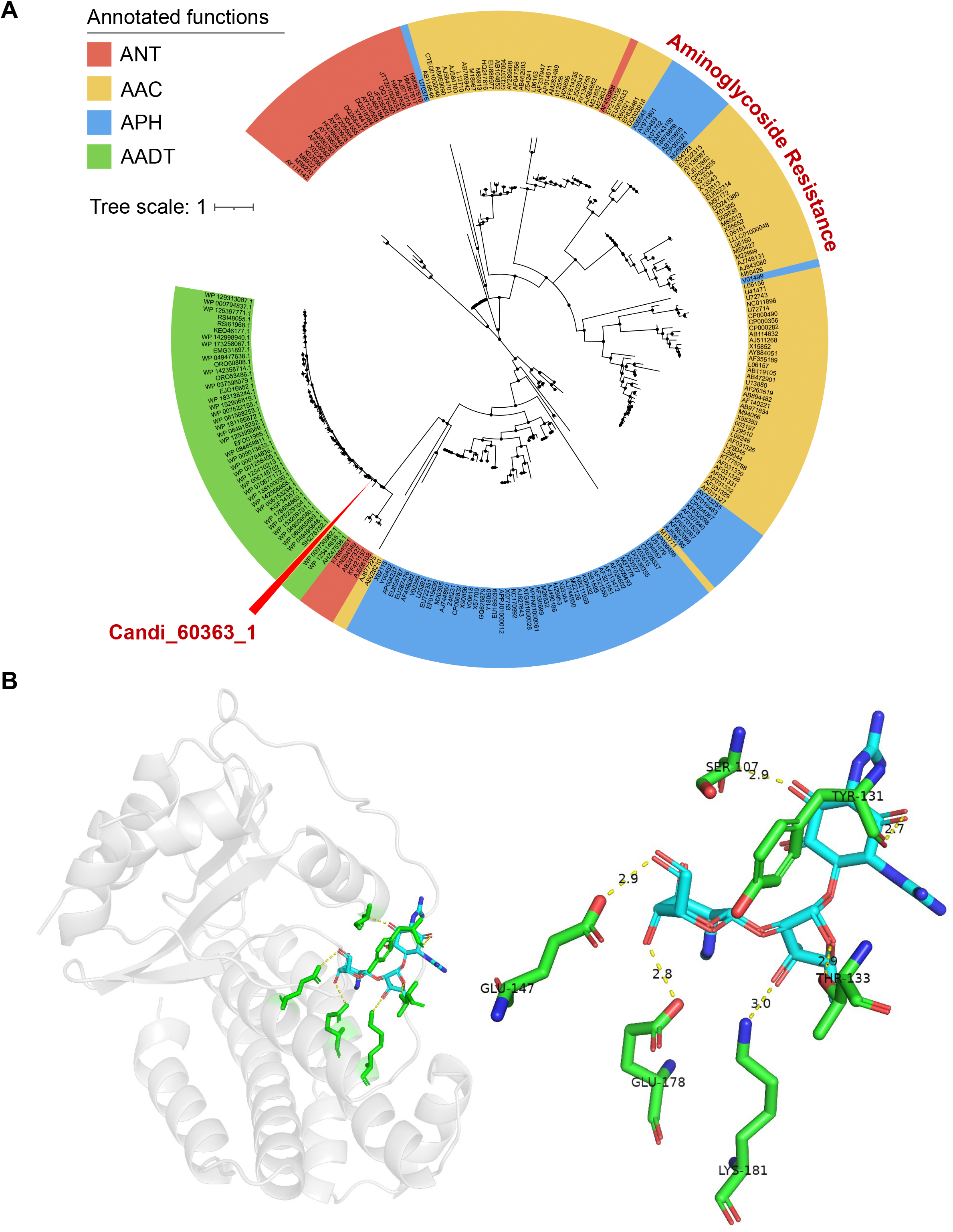
Phylogenetic analysis and structure investigation of Candi_60363_1. (**A**) Phylogenetic tree of aminoglycoside resistance enzymes, Candi_60363_1, and its homologs from the NCBI non-redundant protein database. ANT: O-nucleotidyltransferase, AAC: N-acetyltransferase, APH: O-phosphotransferase, AADT: aminoglycoside adenylyltransferase. (**B**) The optimal Candi_60363_1-streptomycin complex structure (left), and the local interactions between ligand and neighboring residues (right). The docking experiment indicates there are six neighboring residues whose distances are less than three angstroms.

Protein structure prediction results confirmed the anti-microbial functionality of Candi_60363_1. The optimal Candi_60363_1-streptomycin complex structure and the corresponding interaction details are described in **Figure 6B**. The optimal binding affinity between the Candi_60363_1 and streptomycin is −7.7 kcal/mol (**Supplementary Table S11**), which is 1.6 kcal/mol lower than the negative control. From wet-lab experiments, phylogenetic analysis, and protein structure docking, we consider that Candi_60363_1 predicted by ONN4ARG is highly likely a real ARG gene.

## Discussion

In this study, we proposed an ontology-aware deep learning method, ONN4ARG, for the detection and understanding of ARGs. To complement ONN4ARG for ARG mining applications, we have also created a custom ARG database, ONN4ARG-DB, that contains 28,396 well-curated ARGs. The application of ONN4ARG uncovered 120,726 ARGs from microbiome samples, out of which 42,848 are novel, which substantially expands the existing ARGs repositories.

The novelty of this work is in three contexts. First, ONN4ARG has the potential for detection of remotely homologous ARGs and thus generates a more comprehensive set of ARGs. The ability of ONN4ARG to identify remotely homologs allows more accurate prediction. The antibiotic resistance ontology used in the ONN4ARG model consists of four levels and more than 100 resistance subtypes (i.e., terms in the most informative level on the ontology), which substantially expand the classification space of current tools (e.g., 30 types supported for DeepARG and 15 types supported for HMD-ARG). Therefore, ONN4ARG greatly reduces false negatives and offers a powerful approach for accurate and comprehensive profiling of ARGs.

Second, it enabled the comprehensive enrichment analysis of ARGs, species-wise and environment-wise. The environment-specific and host-specific enrichment of ARGs may be caused by specific bacteria evolving to possess specific types of ARGs in response to specific environments, and horizontal gene transfer may be one of the mediating pathways of this process. For example, one published study has reported that *Amycolatopsis* in the soil environment produces rifamycin and thus gains ecological advantages over other bacteria [32].

Third, our study demonstrates the importance and potential of complementing the computational work with wet-lab experimental validation of gene function. Functional verification of a novel streptomycin resistance gene (i.e., Candi_60363_1) with wet-lab experiments demonstrated the ability of the ONN4ARG model for novel ARG discovery. Moreover, phylogenetic analysis and protein structure docking further confirmed that Candi_60363_1 is highly likely to be an ARG that confers resistance to aminoglycoside antibiotics. Another validation of a recently annotated ARG (i.e., GAR) also indicated the ability of the ONN4ARG model for novel ARG discovery.

## Conclusions

We proposed an ontology-aware deep learning approach, ONN4ARG, which is superior to existing methods such as DeepARG in efficiency, accuracy, and comprehensiveness. It enables comprehensive ARG discovery. It has detected novel ARGs that are remotely homologous to existing ARGs. Whereas ONN4ARG has provided one of the most comprehensive profiles of ARGs, it could be further optimized. For more comprehensive ARG prediction, continuous improvement of curating ARG nomenclature and annotation databases is required. For novel ARG prediction, especially those belonging to entirely new ARG families, deep learning models might need to consider more information other than sequence alone, such as protein structure. We believe these efforts could lead to a holistic view about ARGs in diverse environments around the globe.

## Methods

### Dataset

The ARGs we used in this study for model training and testing were from the Comprehensive Antibiotic Resistance Database, CARD v3.0.3 [5, 6]. We also used protein sequences from the UniProt (SwissProt and TrEMBL) database to expand our training dataset. First, genes with ARG annotations were collected from CARD (2,587 ARGs) and SwissProt (2,261 ARGs). Then, their close homologs (sequence identity > 90% and coverage > 98%) were collected from TrEMBL (23,728 homologous genes). These annotated and homologous ARGs made up our ARG dataset. The non-ARG dataset was made from non-ARG genes that had relatively low sequence similarities to ARG genes (sequence identity < 90% and bit-scores < alignment lengths) but not annotated as ARG genes in SwissProt (17,937 non-ARG genes). Finally, redundant genes with identical sequences were filtered out. As a result, our ARG gene dataset, namely, ONN4ARG-DB, contained 28,396 ARG genes and 17,937 non-ARG genes. The gene clustering of the 681 newly added ARGs in CARD v3.1.3 was performed using the MMseqs2 tool (version 10) with an identity of 90% and coverage of 98%. The ResFinder dataset was obtained in Jun 2022 from https://bitbucket.org/genomicepidemiology/resfinder_db/src/master/.

### Antibiotic resistance ontology

The antibiotic resistance ontology was organized into an ontology structure, which contains four levels (**Figure 1A**). The root (first level) is a single node, namely, “arg” (**Supplementary Table S12)**. There are 1, 2, 34, and 277 nodes from the first level to the fourth level, respectively. For instance, there are “beta-lactam” and “non-beta-lactam” in the second level, “acridine dye” and “aminocoumarin” in the third level, and “acriflavine” and “clorobiocin” in the fourth level.

### Framework of ONN4ARG

#### ONN4ARG model

Considering a query gene *q* represented by its protein sequence, as well as its potential resistance categories represented by the antibiotic resistance ontology *O*, to predict resistance categories 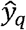 of query gene *q*, we employed ontology-aware neural network to learn a mapping *M* from a set of base genes *b ∈ S* to their resistance categories 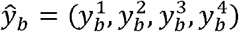. Here, *S* is the set of base genes (i.e., ONN4ARG-DB), 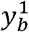 is the resistance category for base gene *b* in the first level of the antibiotic resistance ontology. Then, we apply *M* on *q* to determine the potential resistance categories of query gene.

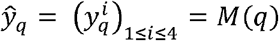

#### Feature encoding

The task of feature encoding is to abstract the homologous signal of a query gene. ONN4ARG takes homologous signals (e.g., identity, e-value, bit-score) between the protein sequence of query gene and protein sequences and profiles (i.e., position-specific scoring matrix) of base genes as features. The homologous signal abstraction works as following. First, a protein sequence library of base genes was made by using “makedb” function of Diamond software. Then, protein sequences of query genes and base genes were aligned by using “blastp” function of Diamond program (**Figure 1B**). Second, profile hidden Markov models (HMMs) of base genes were generated by using “HHblits” function of HH-suite3 software (version 3.2.0). Then, protein sequence of query genes and profile HMMs of base genes were aligned by using “HHblits” function of HH-suite3 software (**Figure 1B**). Third, these homologous signals were normalized (i.e., divided by alignment length) and saved as vectors. The vector sizes at the two-layers of feature embedding network are decided based on the number of sequences and profiles in the ONN4ARG-DB. The vector size of the sequence features is 25,868, and 9,564 for the profile HMMs features.

#### Architecture of the ontology-aware neural network

PyTorch version 1.7.1 was used for generating the ONN model. The architecture of the ontology-aware neural network could be described in four functional layers, including feature embedding layer, residual layer, compress layer and ontology-aware layer (**Supplementary Figure S1**). Details about the four functional layers are available at **Supplementary File S1**.

#### Training and testing

We performed 4-fold cross-validation in the systematic evaluation of ONN4ARG model. In each fold, we divided the ONN4ARG-DB into training set and testing set, the training set contains 75% randomly selected genes from the ONN4ARG-DB, whereas the remaining 25% genes were selected as testing set. We create binary label vector for each protein sequence. If a protein sequence is annotated with a resistance type from the ontology, then we assign 1 to the type’s position in the binary label vector. Otherwise, we assign 0.

#### Masking threshold

To simulate remotely homologous ARG genes in our experiments, homologous signals between the query protein and its close homologs with sequence identities greater than a threshold were masked as zeros (i.e., no signals). For instance, when the masking threshold of testing set equaled 0.4, homologous signals between the query protein (in the testing set) and its close homologs (in the training set) with sequence identities greater than 40% were masked as zeros. Occasionally, all homologs were masked for a query protein, and such query proteins were removed during testing (**Table 1**). For example, if query *X* had two homologs, *M* and *N*, and assuming the identity of *M* is 0.45 and the identity of *N* is 0.95, when the masking threshold of the testing set equaled 0.9, homologous signals between query *X* and homolog *N* were masked as zeros. When the masking threshold of the testing set equaled 0.4, query *X* was removed during testing (see **Table 1** for details).

#### Other methods

We used Diamond (version 0.9.0) [23] as the sequence-alignment tool for comparison. We used the same training and testing sets as in the ONN4ARG model to evaluate the sequence-alignment method. For queries in the testing set, we searched them against the training set. The target with the highest identity was defined as the closest homologous gene for each query. Then, we compared whether the actual annotation of the query was consistent with the annotation of its closest homologous gene to evaluate the performance. DeepARG [12] is a newly developed tool that applies a plain neural network (e.g., several fully connected layers) to predict ARGs. Here, we reconstructed the DeepARG model with PyTorch by using the same architecture of original DeepARG model, and used the same training and testing sets as in the ONN4ARG model to train and test the DeepARG model. For queries in the testing set, we used the reconstructed DeepARG model to predict their ARG annotations, and compared whether the actual annotations were consistent with the predicted annotations to evaluate the performance.

#### Performance measures

To assess the performance of ONN4ARG model and other methods, we used accuracy measure with the following formula:

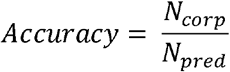

where *N_corp_* is the number of correct predictions, and *N_pred_* is the number of total predictions. Notably, a prediction was defined to be correct if and only if all ARG annotations (including ancestor annotations from ARG ontology) were correctly predicted.

Furthermore, we used precision, recall, F1, AUROC, and AUPRC measures to assess the performance of ONN4ARG model and other methods on each antibiotic resistance type:

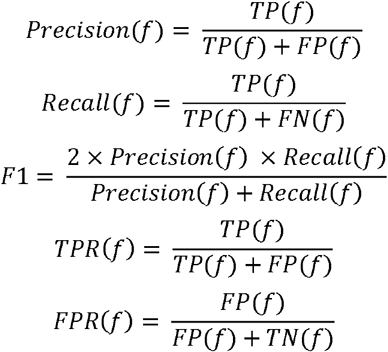

where *f* represents one resistance type, *TP*(*f*) is the number of true positive predictions of resistance type *f, FP*(*f*) is the number of false positive predictions of resistance type *f, TN*(*f*) is the number of true negative predictions of resistance type *f*, and *FN*(*f*) is the number of false negative predictions of resistance type *f*. AUROC is the area under the *TPR-FPR* curve, and AUPRC is the area under the *Precision-Recall* curve.

#### Taxonomy annotation

Kraken2 (version 2.1.2) [29] program with default parameters was used to identify the host of gene contigs. Then, each ARG predicted by ONN4ARG was annotated according to the host of its gene contigs.

#### Phylogenetic tree

Protein sequences of the most closely related to Candi_60363_1 were collected using BLASTP with default parameters on the NCBI non-redundant protein database. The retrieved proteins, Candi_60363_1 and all aminoglycoside resistance proteins from ResFinder [8] (https://bitbucket.org/genomicepidemiology/resfinder_db/src/master, last update Jun 2022), were aligned with ClustalW. The phylogenetic tree was calculated by MEGA [39] (v10) using the maximum likelihood algorithm with default parameters. The Interactive Tree of Life (iTOL v6) online tool [40] was used to prepare the phylogenetic tree for display.

#### Protein model and docking

Rosetta [41] was utilized to predict the protein structure using ab initio protein folding (http://robetta.bakerlab.org/). The top five protein pockets were generated for docking calculation with Surface Topography of proteins [42] (CASTp). We used the Cambridge Structure Database [43] to generate streptomycin conformers. The 3D protein-ligand complexes were obtained from AutoDock Vina [44].

#### ARG candidate gene expression plasmids construction and expression verification

The candidate resistance gene Candi_60363_1 and a positive control resistance gene AHE40557.1 were synthesized and subcloned into pUC19 vector, replacing *lacZ’* gene. The recombinant plasmids were then transformed into *E. coli* BL21 (DE3). The expression of resistance genes was induced by Isopropyl β-D-1-thiogalactopyranoside (IPTG) and verified by quantitive Real-time PCR (qRT-PCR) assay. Briefly, bacteria were grown in LB supplemented with ampicillin (100 μg/ml) to OD600 of 0.5-0.6 by incubation at 37 °C with 220 rpm agitation, and the bacterial cultures were continued to grow until OD600 reached to 1.0 by adding or without adding 1 mM IPTG. The cells were harvested and total RNAs were purified using Bacterial RNA Extraction Kit (Vazyme Biotech). RNA reverse transcription was performed by using HiScript® II Q Select RT SuperMix for qPCR kit (Vazyme Biotech). qRT-PCR was performed by using SYBR Green Master Mix-High ROX Premixed (Vazyme Biotech) in a Stepone Plus system (Applied Biosystems). The *ldh* gene was used as internal control in all reactions. The relative fold changes were determined using the 2^-ΔΔCt^ method, in which *ldh* was used for normalization. The protein sequences of the synthesized genes and the primer sequences for qRT-PCR were listed in **Supplementary Table S8** and **S9**.

#### MIC determination

Minimal inhibitory concentrations (MICs) of the antibiotic for the strains containing resistance genes were determined using E-tests (three repeats). Single colonies of the strains were incubated in 3 ml Mueller-Hinton (MH) medium with the addition of 100 μg/ml ampicillin at 35 ^o^C for 4 hours, and the cells equal to 1.5X10^8^ cells/ml were spread on MH agar plates with the addition of 100 μg/ml ampicillin and 1 mM IPTG, and streptomycin MIC Test Strips (Liofilchem®) were put in the middle of the plates. The plates were incubated at 35 ^o^C for 18-24 hours, and the MICs were read. The strain containing empty vector was used as a negative control.

#### Statistical test

According to the normality of the data distribution verified by the Shapiro–Wilk test and Levene’s test, the ARG abundance data distribution is Gaussian and unequal variance. Thus, statistical test of the enrichment analysis was performed utilizing the Welch’s *t*-test (one-tailed), at the significance level of 0.005 [45]. For all the tests, when the *P* value associated is lower than the significance level, one should reject the null hypothesis H0 (ARGs are not enriched in the environment or host), and accept the alternative hypothesis Ha (ARGs are enriched in the environment or host).

## Key Points

- We developed an ontology-aware deep learning approach, ONN4ARG, which is superior to existing methods such as DeepARG in efficiency, accuracy.
- ONN4ARG has the potential for detection of remotely homologous ARGs and thus generates a more comprehensive set of ARGs.
- ONN4ARG enabled the comprehensive enrichment analysis of ARGs, species-wise and environment-wise.
- Our study demonstrates the importance and potential of complementing the computational work with wet-lab experimental validation of gene function.

## Declarations

### Ethics approval and consent to participate

Not applicable

### Consent for publication

Not applicable

### Competing interests

The authors declare that they have no competing interests.

### Data availability

We collected metagenomic samples from several published studies [25, 26], and these samples are mainly from marine, soil and human associated environments. For human associated samples, including two gut groups (one group from Madagascar, i.e., GutM, the other group from Denmark, i.e., GutD), one oral group and one skin group (both oral and skin groups are from HMP project). Details and links about these samples are shown in **Supplementary Table S3**. The ONN4ARG-DB dataset could be accesses at: https://github.com/HUST-NingKang-Lab/ONN4ARG.

### Code availability

All source codes have been uploaded to the website at: https://github.com/HUST-NingKang-Lab/ONN4ARG, and online web service can be accessed at: http://onn4arg.xfcui.com/.

### Authors’ contributions

K.N, X.C conceived and proposed the idea, and designed the study. Y.Z, C.C, Q.J, X.Z, X.C performed the experiments and analyzed the data. Y.Z, C.C, X.Z, K.N and X.C contributed to editing and proof-reading the manuscript. All authors read and approved the final manuscript.

## Acknowledgments

We are grateful to Mingyue Cheng for insightful discussions. This work was partially supported by National Natural Science Foundation of China (Grant Nos. 81774008, 81573702, 31871334 and 31671374), and the National Key R&D Program (Grant No. 2018YFC0910502).

**Yuguo Zha** is a PhD student at the Huazhong University of Science and Technology. His research interests are in microbiome associated data mining, including gene mining, species mining, and pattern mining.

**Cheng Chen** is a PhD student at Shandong University. His research interests are in computational biology and bioinformatics.

**Qihong Jiao** is a PhD student at Shandong University. His research interests are in computational biology and bioinformatics.

**Xiaomei Zeng** is a professor at Department of Bioinformatics and Systems Biology, College of Life Science and Technology, Huazhong University of Science and Technology.

**Xuefeng Cui** is a professor at the School of Computer Science and Technology, Shandong University. His research interests are in computational biology and bioinformatics.

**Kang Ning** is a professor at Department of Bioinformatics and Systems Biology, College of Life Science and Technology, Huazhong University of Science and Technology.

---

**Table 1. Accuracy comparison of sequence-alignment, DeepARG and ONN4ARG based on different masking threshold of testing set.**

**Table 2. Evaluation results of ONN4ARG for ARGs identification at different masking threshold of testing set.**

**Table 3. Evaluation results of DeepARG for ARGs identification at different masking threshold of testing set.**

## Supplementary Materials

**Supplementary Figure S1. The architecture of the ontology-aware neural network.** (**A**) The architecture of the ontology-aware neural network could be described in four functional layers, including feature embedding layer, residual layer, compress layer and ontology-aware layer. The ontology-aware layer is a partially connected layer which encourage annotation predictions satisfying the ontology rules (i.e., the ontology tree structure). Specially, weight between nodes with relationship (e.g., parent and child) satisfying the ontology rules would be saved in the partially connected layer, and weights between irrelevant nodes would be masked. (**B**) The weight matrix derived from the antibiotic resistance ontology and the ontology-aware layer.

**Supplementary Figure S2. The number of pan and core ARG types among various environments, and gene mobility analysis for predicted ARGs.** (**A**) The number of pan and core ARG types change as more groups are included. For core/pan counts, we only counted ARG types with the relative abundance ratio greater than 1e-4. The pan ARGs refer to the ARG types that are included in any environments. The core ARGs refer to the ARG types that are included in all environments. (**B**) The venn diagram shows the ARG types relationship among marine, soil and gut groups. (**C**) The venn diagram shows the ARG types relationship among gut, oral and skin groups. (**D**) The distribution of acquired and intrinsic ARGs in various environments. (**E**) The line regression analysis indicates no significant correlation (*P* > 0.05) between the abundances of MGEs and ARGs. The horizontal axis indicates the abundance ratio of predicted ARGs and the vertical axis indicates the abundance ratio of MGEs. Each point represents a group.

**Supplementary Figure S3. The host range of all classified ARGs and the resistance composition of the most abundant 20 genera.** (**A**) The Sankey diagram shows the host composition and distribution of all classified ARGs (the most abundant 20 genera carrying ARGs were used for display). (**B**) The bar chart indicates the diversity and relative abundance of ARGs for the most abundant 20 genera carrying ARGs.

**Supplementary Figure S4. The network analysis revealing the co-occurrence patterns among ARG types and microbial taxa, the nodes were represented by pie charts which shows the taxonomic compositions of ARG types.** A connection represents a strong (Spearman’s *ρ* > 0.8) and significant (*P*-value < 0.01) correlation. The size of each node is proportional to the number of connections, that is, the degree.

**Supplementary Figure S5. The MIC experiment for predicted candidate ARG (top), negative control (middle) and positive control (bottom).** The MIC values are tested for three repeats.

**Supplementary Table S1. Comparison of ONN4ARG and other methods for ARG identification on the verification set.**

**Supplementary Table S2. Evaluation of ONN4ARG and DeepARG on the ResFinder dataset.**

**Supplementary Table S3. Metagenomic samples using for resistance gene mining are collected from published studies.**

**Supplementary Table S4. The number of predicted ARGs by ONN4ARG that have protein domains with known catalytic activity and/or may bind to the antimicrobials they are predicted to elicit resistance against.**

**Supplementary Table S5. Data distribution during the pipeline of ARGs prediction.**

**Supplementary Table S6. The hosts of predicted ARGs at different taxonomic level.**

**Supplementary Table S7. The predicted ARGs which fulfilling all mentioned criteria.**

**Supplementary Table S8. Protein sequences of the synthesized genes.**

**Supplementary Table S9. Real-time PCR primer sequences.**

**Supplementary Table S10. The BLAST result of Candi_60363_1 when search against the NCBI non-redundant protein database.**

**Supplementary Table S11. The binding affinity of protein–ligand complexes using the top five pockets.**

**Supplementary Table S12. The antibiotic resistance ontology used in the ONN4ARG model.**

**Supplementary File S1. Supplemental information about experiments.**

## Notes

### Competing Interest Statement

The authors have declared no competing interest.

### Summary of Updates

In this revision, we have fixed some typos and grammar errors. We also added some experiments and new results in the revision.

## References

1. Thomas AM, Segata N. Multiple levels of the unknown in microbiome research. BMC Biol. 2019;17:48.

2. Li J, Jia H, Cai X, Zhong H, Feng Q, Sunagawa S, et al. An integrated catalog of reference genes in the human gut microbiome. Nat Biotechnol. 2014; 32:834–841.

3. Brogan DM, Mossialos E. A critical analysis of the review on antimicrobial resistance report and the infectious disease financing facility. Global and Health. 2016;12:8.

4. Goossens H, Ferech M, Stichele RV, Elseviers M. Outpatient antibiotic use in Europe and association with resistance: a cross-national database study. Lancet 2005;365:579–587.

5. Jia B, Raphenya AR, Alcock B, Waglechner N, Guo P, Tsang KK, et al. CARD 2017: expansion and model-centric curation of the comprehensive antibiotic resistance database. Nucleic Acids Res. 2017;45:D566–D573.

6. Alcock BP, Raphenya AR, Lau TTY, Tsang KK, Bouchard M, Edalatmand A, et al. CARD 2020: antibiotic resistome surveillance with the comprehensive antibiotic resistance database. Nucleic Acids Res. 2020;48:D517–D525.

7. Böhm M-E, Razavi M, Marathe NP, Flach C-F, Larsson DGJ. Discovery of a novel integron-borne aminoglycoside resistance gene present in clinical pathogens by screening environmental bacterial communities. Microbiome 2020;8:1–11.

8. Bortolaia V, Kaas RS, Ruppe E, Roberts MC, Schwarz S, Cattoir V, et al. ResFinder 4.0 for predictions of phenotypes from genotypes. J Antimicrob Chemother. 2020;75:3491–3500.

9. Bateman A, Martin MJ, O’Donovan C, Magrane M, Apweiler R, Alpi E, et al. UniProt: A hub for protein information. Nucleic Acids Res. 2015;43: D204–D212.

10. Rowe W, Baker KS, Verner-Jeffreys D, Baker-Austin C, Ryan JJ, Maskell DJ, et al. Search Engine for Antimicrobial Resistance: a cloud compatible pipeline and web interface for rapidly detecting antimicrobial resistance genes directly from sequence data. PLoS One. 2015;10:e0133492.

11. Kleinheinz KA, Joensen KG, Larsen MV. Applying the ResFinder and VirulenceFinder web-services for easy identification of acquired antibiotic resistance and E. coli virulence genes in bacteriophage and prophage nucleotide sequences. Bacteriophage. 2014;4:e27943.

12. Arango-Argoty G, Garner E, Pruden A, Heath LS, Vikesland PJ, Zhang L. DeepARG: a deep learning approach for predicting antibiotic resistance genes from metagenomic data. Microbiome. 2018;6:23.

13. Davis JJ, Boisvert S, Brettin T, Kenyon RW, Mao C, Olson R, et al. Antimicrobial resistance prediction in PATRIC and RAST. Sci Rep. 2016;6:27930.

14. Lakin SM, Kuhnle A, Alipanahi B, Noyes NR, Dean C, Muggli M, et al. Hierarchical Hidden Markov models enable accurate and diverse detection of antimicrobial resistance sequences. Commun Biol. 2019;2:294.

15. Doster E, Lakin SM, Dean CJ, Wolfe C, Young JG, Boucher C, et al. MEGARes 2.0: a database for classification of antimicrobial drug, biocide and metal resistance determinants in metagenomic sequence data. Nucleic Acids Res. 2020;48:D561–D569.

16. Li Y, Xu Z, Han W, Cao H, Umarov R, Yan A, et al. HMD-ARG: hierarchical multi-task deep learning for annotating antibiotic resistance genes. Microbiome 2021;9:40.

17. Gupta SK, Padmanabhan BR, Diene SM, Lopez-Rojas R, Kempf M, Landraud L, et al. ARG-ANNOT, a new bioinformatic tool to discover antibiotic resistance genes in bacterial genomes. Antimicrob Agents Chemother. 2014;58:212–220.

18. Feldgarden M, Brover V, Haft DH, Prasad AB, Slotta DJ, Tolstoy I, et al. Validating the AMRFinder tool and resistance gene database by using antimicrobial resistance genotype-phenotype correlations in a collection of isolates. Antimicrob Agents Chemother. 2019;63:e00483.

19. Inouye M, Dashnow H, Raven LA, Schultz MB, Pope BJ, Tomita T, et al. SRST2: rapid genomic surveillance for public health and hospital microbiology labs. Genome Med. 2014;6:90.

20. Rowe WPM, Winn MD. Indexed variation graphs for efficient and accurate resistome profiling. Bioinformatics. 2018;34:3601–3608.

21. Altschul SF, Gish W, Miller WC, Myers EW, Lipman DJ. Basic Local Alignment Search Tool. J Mol Biol. 1990;215:403–410.

22. Edgar RC. Search and clustering orders of magnitude faster than BLAST. Bioinformatics. 2010;26:2460–2461.

23. Buchfink B, Xie C, Huson DH. Fast and sensitive protein alignment using DIAMOND. Nat Methods. 2015;12:59–60.

24. Steinegger M, Meier M, Mirdita M, Vöhringer H, Haunsberger SJ, Söding J. HH-suite3 for fast remote homology detection and deep protein annotation. BMC Bioinform. 2019;20:1–15.

25. Sunagawa S, Coelho LP, Chaffron S, Kultima JR, Labadie K, Salazar G, et al. Structure and function of the global ocean microbiome. Science. 2015;348:1261359.

26. Mitchell AL, Scheremetjew M, Denise H, Potter S, Tarkowska A, Qureshi M, et al. EBI Metagenomics in 2017: enriching the analysis of microbial communities, from sequence reads to assemblies. Nucleic Acids Res. 2018;46:D726–D735.

27. Hyatt D, Chen GL, LoCascio PF, Land ML, Larimer FW, Hauser LJ. Prodigal: prokaryotic gene recognition and translation initiation site identification. BMC Bioinform. 2010;11:119.

28. Li B, Yang Y, Ma L, Ju F, Guo F, Tiedje JM, et al. Metagenomic and network analysis reveal wide distribution and co-occurrence of environmental antibiotic resistance genes. ISME J. 2015;9:2490–2502.

29. Wood DE, Lu J, Langmead B. Improved metagenomic analysis with Kraken 2. Genome Biol. 2019;20:1–13.

30. Qi F, Lei C, Li F, Zhang X, Wang J, Zhang W, et al. Deciphering the late steps of rifamycin biosynthesis. Nat Commun. 2018;9:2342.

31. Floss HG, Yu T-W. Rifamycin-mode of action, resistance, and biosynthesis. Chem Rev. 2005;105:621–632.

32. Yao Y, Zhang W, Jiao R, Zhao G, Jiang W. Efficient isolation of total RNA from antibiotic-producing bacterium Amycolatopsis mediterranei. J Microbiol Methods. 2002;51:191–195.

33. Wilson MC, Gulder TAM, Mahmud T, Moore BS. Shared biosynthesis of the saliniketals and rifamycins in Salinispora arenicola is controlled by the sare1259-encoded cytochrome P450. J Am Chem Soc. 2010;132:12757–12765.

34. Saxena A, Kumari R, Mukherjee U, Singh P, Lal R. Draft genome sequence of the rifamycin producer amycolatopsis rifamycinica DSM 46095. Genome Announc. 2014;2:e00662.

35. Huang H, Lv J, Hu Y, Fang Z, Zhang K, Bao S. Micromonospora rifamycinica sp. nov., a novel actinomycete from mangrove sediment. Int J Syst Evol Microbiol. 2008;58:17–20.

36. Pinto-Alphandary H, Mabilat C, Courvalin P. Emergence of aminoglycoside resistance genes aadA and aadE in the genus Campylobacter. Antimicrob Agents Chemother. 1990;34:1294–1296.

37. Holden MTG, Hauser H, Sanders M, Ngo TH, Cherevach I, Cronin A, et al. Rapid evolution of virulence and drug resistance in the emerging zoonotic pathogen Streptococcus suis. PLoS One. 2009;4:e6072.

38. Ramirez MS, Nikolaidis N, Tolmasky M. Rise and dissemination of aminoglycoside resistance: the aac(6’)-Ib paradigm. Front Microbiol. 2013;4:121.

39. Kumar S, Stecher G, Li M, Knyaz C, Tamura K. MEGA X: molecular evolutionary genetics analysis across computing platforms. Mol Biol Evol. 2018;35:1547–1549.

40. Letunic I, Bork P. Interactive Tree Of Life (iTOL) v4: recent updates and new developments. Nucleic Acids Res. 2019;47:W256–W259.

41. Rohl CA, Strauss CEM, Misura KMS, Baker D. Protein structure prediction using Rosetta. Methods Enzymol. 2004;383:66–93.

42. Tian W, Chen C, Lei X, Zhao J, Liang J. CASTp 3.0: Computed Atlas of Surface Topography of Proteins. Nucleic Acids Res. 2018;46:W363–W367.

43. Cole JC, Korb O, McCabe P, Read MG, Taylor R. Knowledge-based conformer generation using the cambridge structural database. J Chem Inf Model. 2018;58:615–629.

44. Trott O, Olson AJ. AutoDock Vina: improving the speed and accuracy of docking with a new scoring function, efficient optimization, and multithreading. J Comput Chem. 2010;31:455–461.

45. Benjamin DJ, Berger JO, Johannesson M, Nosek BA, Wagenmakers EJ, Berk R, et al. Redefine statistical significance. Nat Hum Behav. 2018;2:6–10.

