## Supplementary material for "Ontology-aware deep learning for antibiotic resistance gene prediction: novel function discovery and comprehensive profiling from metagenomic data": Supplementary Figure S5.pdf

### Candi_60363_1

| Repeat 1 | Repeat 2 | Repeat 3 |
|---|---|---|
| 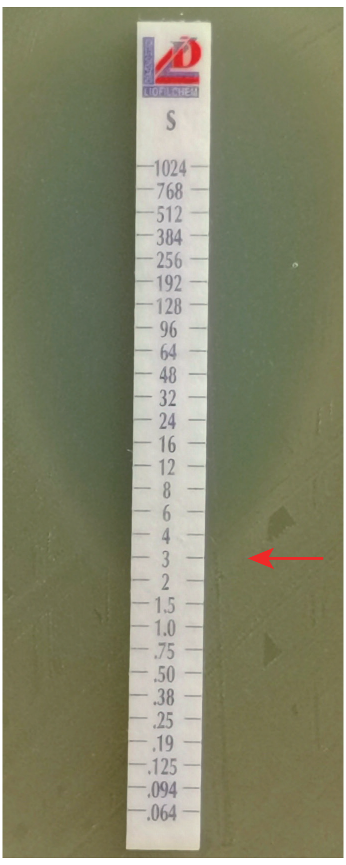 | 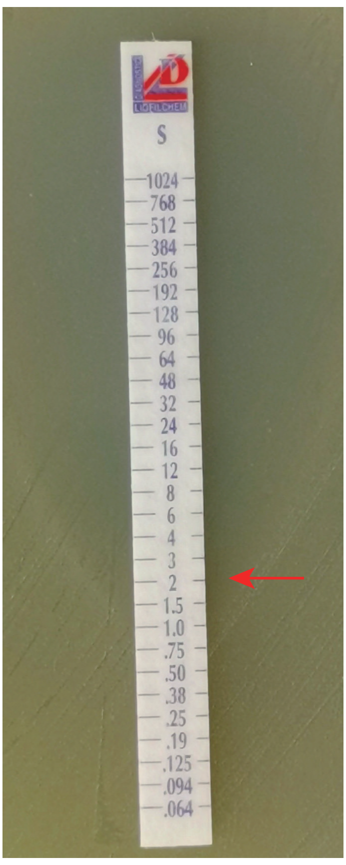 | 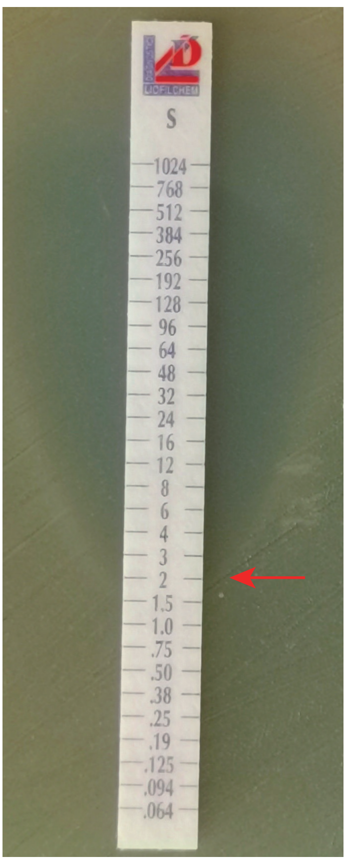 |
| MIC = 3 | MIC = 2 | MIC = 2 |

### Negative control

| Repeat 1 | Repeat 2 | Repeat 3 |
|---|---|---|
| 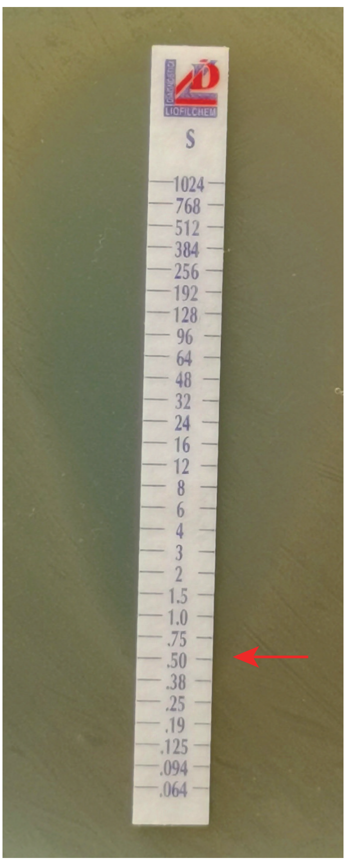 | 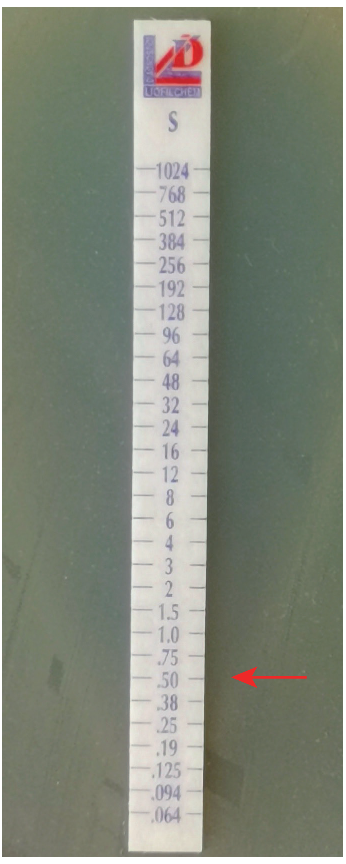 | 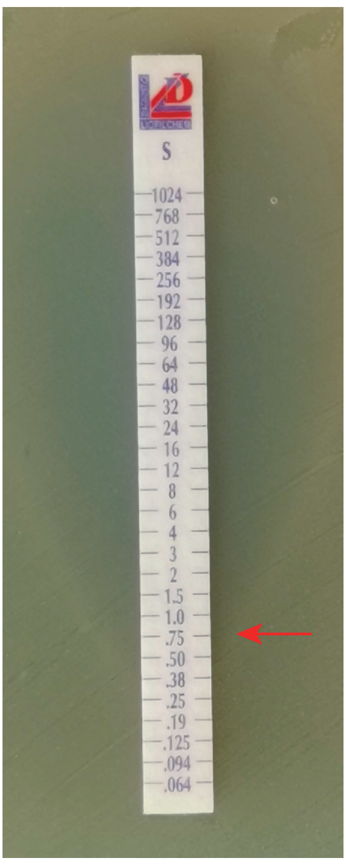 |
| MIC = 0.5 | MIC = 0.5 | MIC = 0.75 |

### AHE40557.1

| Repeat 1 | Repeat 2 | Repeat 3 |
|---|---|---|
| 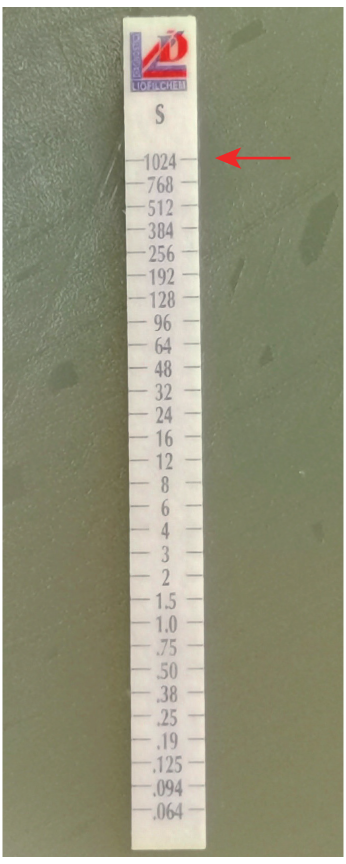 | 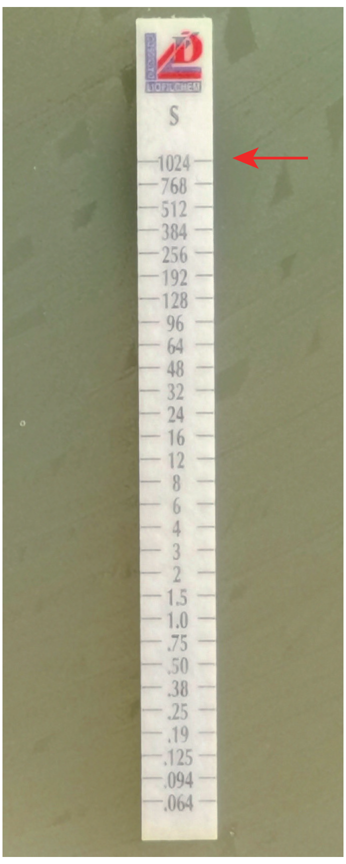 | 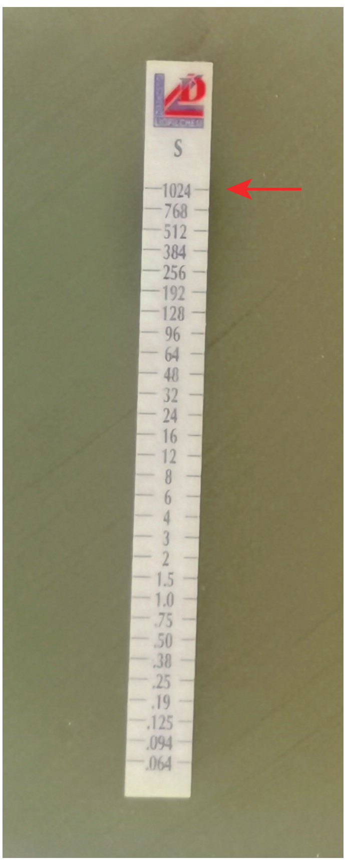 |
| MIC > 1024 | MIC > 1024 | MIC > 1024 |
