## Supplementary material for "Ontology-aware deep learning for antibiotic resistance gene prediction: novel function discovery and comprehensive profiling from metagenomic data": Supplementary File S1.docx

**Supplementary Note: *The generalization and data augmentation experiments of ONN4ARG***

One desired property of any machine learning model is to have good generalization abilities to unseen data with different properties and distributions. For example, after training the model for mostly close homologs (which is usually the case), we would like the model to predict accurate annotations for not only close but also remote homologs. To evaluate such model generalization abilities, similarities between the query protein and its close homologs with sequence identities greater than a threshold are masked as zeros (i.e., no signals), and different combinations of training and testing masking thresholds are tested in **Table 1**. Occasionally, all homologs are masked for a query protein, and such query proteins are removed during training and testing.

Due to the outstanding generalization abilities, ONN4ARG achieves the highest average accuracies among all tested methods (**Fig. 2**). Interestingly, sequence alignment consistently performs reasonably well for close homologs, but this is not the case for remote homologs. That suggests that sequence alignment is less sensitive to the differences between training and testing datasets. One possible explanation is that its predictions are only related to the closest homolog instead of all homologs used by deep learning methods. Although ONN4ARG is shown to have the highest accuracies and the best generalization abilities in this experiment, it is still a good idea to train a model with both close and remote homologs. Thus, all masked training data in this experiment is combined as a bigger dataset, and another ONN4ARG model is retrained with the new dataset for the following experiments. This step can be seen as a data augmentation technique which usually produces higher accuracies (**Tables 2** and **3**).

**Supplementary Note:** ***Co-occurrence patterns among ARG types and microbial taxa***

The co-occurrence patterns among ARG types were explored using network inference based on strong (Pearson’s correlation $\rho$ > 0.8) and significant (*P*-value < 0.01) correlations. The interrelationships among the ARG types which belong to beta-lactam resistance type (i.e., cephamycin, penam, penem, and monobactam) were clearly observed. One possible explanation for the interrelationships among ARG types belonging to β-lactams category is that they might be harbored in some specific microbial taxa that are shared by different environments. In addition, these ARGs types belonging to β-lactams are likely to appear together in some specific microbial taxa since β-lactams antibiotics have similar bactericidal mechanism.

To understand the host compositions of ARG types in the co-occurrence network, we used pie charts to show the host contributions for all the nodes. Results showed that Actinobacteria and Proteobacteria are the most abundant phyla (70 % on average, **Supplementary Fig. 4**) in all the ARG types. The non-random co-occurrence patterns between ARGs and microbial taxa could indicate the possible host information of ARGs if the ARGs and the co-existed microbial taxa possessed the significantly similar abundance trends among the different environments (Spearman’s $\rho$ > 0.8, P-value < 0.01).

**Supplementary Note: *Details about the four functional layers in the ONN***

The feature embedding layer comprises two fully connected layers. The first fully connected layer accepts the flattened features (e.g., sequence alignment features) and outputs a vector of size 4,096. The input vector sizes are decided based on the number of sequences and profiles in the ONN4ARG-DB. The vector size of the sequence features is 25,868, and 9,564 for the profile HMMs features. Then, the output vector is operated with “Group Norm” and “GELU” functions and passed to the second fully connected layer. The output of the second fully connected layer is an embedding vector of size 1,024.

The residual layer comprises four fully connected layers (with “Group Norm” and “GELU” functions), which learn residual functions with reference to the layer inputs, instead of learning unreferenced functions. Formally, denoting the desired underlying mapping as $H\left( x \right)$, we let the stacked nonlinear layers fit another mapping of $F\left( x \right)=H\left( x \right)-x$. The original mapping is recast into $F\left( x \right)+x$. The $F\left( x \right)$ acts like a residual, hence the name residual layer. By incorporating the residual layer into the ONN4ARG model, the accuracy and generalization ability could be improved and reduce the overfitting.

The compress layer is a simple fully connected layer (with “Group Norm” and “GELU” functions), which is used to change the size of embedding vector. For example, the output embedding vector of residual layer is 1,024, and it would be compressed to the size of 193 (the number of nodes in the antibiotic resistance ontology). Then, the compressed vector is passed to the ontology-aware layer.

The ontology-aware layer is a partially connected layer which encourage annotation predictions satisfying the ontology rules (i.e., the hierarchical structure of the antibiotic resistance ontology). The ontology weight matrix derived from the antibiotic resistance ontology is used to encode the connection between terms at each level of the ontology. Specially, weight between terms with relationship (e.g., parent and child) satisfying the ontology rules would be saved in the partially connected layer, and weights between irrelevant terms would be masked. The ontology-aware layer accepts an embedding vector $V_{pre}$. Then, “Layer Norm” and “GELU” functions will be applied to the vector $V_{pre}$. Finally, the partially connected layer is applied to generate a vector $V_{post}$. The output of ontology-aware layer is $V_{pre}+ V_{post}$.
