## Supplementary material for "Ontology-aware deep learning for antibiotic resistance gene prediction: novel function discovery and comprehensive profiling from metagenomic data": Supplementary Table S1.docx

**Supplementary Table S1. Comparison of ONN4ARG and other methods for ARG identification on the verification set.**

| **Method** | **Accuracy (%) ^a^** | **Time usage (s) ^b^** | **Memory usage (GB) ^c^** |
| --- | --- | --- | --- |
| Sequence-alignment | 44.3 | 2 | 0.1 |
| DeepARG | 56.2 | 10 | 0.6 |
| ONN4ARG | 88.7 | 4 ^d^ | 2.0 |

*Note*: ONN4ARG model was constructed based on the CARD version 3.0.3. We downloaded the new CARD version 3.1.3, which contains 681 new ARGs over the old version 3.0.3. These 681 ARGs were clustered into 203 ARG clusters, and these 203 representative ARGs made up the verification set. DNN, deep neural network; ONN, ontology-aware neural network; ^a^, accuracy is the ratio of correct predictions; ^b^, running on a Linux platform with 20 cores; ^c^, maximum memory usage when programs running; ^d^, query time only (excluding training time).
