## Supplementary material for "Ontology-aware deep learning for antibiotic resistance gene prediction: novel function discovery and comprehensive profiling from metagenomic data": Supplementary Table S2.docx

**Supplementary Table S2. Evaluation of ONN4ARG and DeepARG on the ResFinder dataset.**

| **Resistance** | **No. genes** | **Detailed-classified** | **Identical-classified** | **Under-classified** | **Mis-classified** | **Acc-ONN4ARG** | **Acc-DeepARG** |
| --- | --- | --- | --- | --- | --- | --- | --- |
| Aminoglycoside | 265 | 0.8151 | 0.1321 | 0.0226 | 0.0302 | 0.9472 | 0.8830 |
| Beta-lactam | 2,013 | 0.8107 | 0.1873 | 0 | 0.0020 | 0.9980 | 0.7615 |
| Colistin | 56 | 0 | 1.0000 | 0 | 0 | 1.0000 | 0 |
| Fosfomycin | 41 | 0 | 0.3902 | 0.5854 | 0.0244 | 0.3902 | 0.5122 |
| Glycopeptide | 44 | 0 | 0.9318 | 0.0682 | 0 | 0.9318 | 0.9091 |
| Macrolide | 176 | 0.1250 | 0.1648 | 0.5227 | 0.1875 | 0.2898 | 0 |
| Nitroimidazole | 14 | 0 | 0 | 0 | 1.0000 | 0 | 0 |
| Oxazolidinone | 24 | 0 | 0.1250 | 0 | 0.8750 | 0.1250 | 0 |
| Phenicol | 48 | 0.5208 | 0.3750 | 0.0417 | 0.0625 | 0.8958 | 0.8125 |
| Tetracycline | 148 | 0.4730 | 0.4595 | 0.0203 | 0.0473 | 0.9325 | 0.7973 |
| Trimethoprim | 109 | 0 | 0.9633 | 0.0183 | 0.0183 | 0.9633 | 0 |

***Note***: The ResFinder dataset used in this study is version of 4.1, obtained in Jun 2022. Detailed-classified means the predicted type of resistance by ONN4ARG was more detailed than the actual type of resistance. Identical-classified means the predicted type of resistance by ONN4ARG was the same as the actual type of resistance. Under-classified means the predicted type of resistance by ONN4ARG was coarse. Mis-classified means the predicted type of resistance by ONN4ARG was incorrect. For example, Gar_1_NG_070891 is annotated in the ResFinder dataset as an aminoglycoside resistance gene. The aminoglycoside has a parent of non-beta-lactam and a child of streptomycin in the antibiotic resistance ontology. If the predicted type of Gar_1_NG_070891 is non-beta-lactam/aminoglycoside/streptomycin, then this prediction should be counted as under-classified/identical-classified/detailed-classified. No. genes, number of genes; Acc-ONN4ARG, accuracy of ONN4ARG (detailed-classified plus identical-classified); ACC-DeepARG, accuracy of DeepARG.
