## Supplementary material for "Ontology-aware deep learning for antibiotic resistance gene prediction: novel function discovery and comprehensive profiling from metagenomic data": Supplementary Table S4.docx

**Supplementary Table S4.** **The number of predicted ARGs by ONN4ARG that have protein domains with known catalytic activity and/or may bind to the antimicrobials they are predicted to elicit resistance against.**

|  | Marine | Soil | GutM | GutD | Oral | Skin |
| --- | --- | --- | --- | --- | --- | --- |
| No. ARGs-CDD | 40,899 | 38,467 | 12,914 | 23,981 | 2,467 | 454 |
| No. ARGs | 41,082 | 39,553 | 12,971 | 24,154 | 2,500 | 466 |
| Ratio | 99.56% | 97.25% | 99.56% | 99.28% | 98.68% | 97.42% |

*Note*: No. ARGs, number of predicted ARGs by ONN4ARG; No. ARGs-CDD, number of predicted ARGs by ONN4ARG that have protein domains with known catalytic activity and/or may bind to the antimicrobials they are predicted to elicit resistance against, protein domains are predicted by searching the conserved domain database (CDD) using RPS-BLAST tool; Ratio, No. ARGs-CDD / No. ARGs.
