## Supplementary material for "Ontology-aware deep learning for antibiotic resistance gene prediction: novel function discovery and comprehensive profiling from metagenomic data": Supplementary Table S5.docx

**Supplementary Table S5. Data distribution during the pipeline of ARGs prediction.**

| **Group** | **No. Genes** | **No. CD-HIT** | **No. Diamond** | **No. ONN4ARG** | **No. ARGs** | **No. ARGs / No. Genes** |
| --- | --- | --- | --- | --- | --- | --- |
| Marine | 127,740,693 | 25,489,046 | 1,048,137 | 41,082 | 284,444 | 0.22% |
| Soil | 9,383,234 | 4,155,938 | 215,153 | 39,553 | 98,699 | 1.05% |
| GutM | 10,413,165 | 2,455,995 | 74,358 | 12,971 | 96,618 | 0.93% |
| GutD | 34,222,886 | 4,225,128 | 133,436 | 24,154 | 284,002 | 0.83% |
| Oral | 2,136,190 | 432,790 | 12,495 | 2,500 | 30,578 | **1.43%** |
| Skin | 265,122 | 73,333 | 2,242 | 466 | 2,730 | 1.03% |
| UniProt | 184,998,855 | # | # | # | 579,357 | 0.31% |
| SwissProt | 562,755 | # | # | # | 2,649 | 0.47% |

*Note*: No. Genes, number of genes; No. CD-HIT, number of representative genes after clustering; No. Diamond, number of genes after identity screening; No. ONN4ARG, number of antibiotic resistance genes predicted by ONN4ARG model; No. ARGs, number of resistance genes; No. ARGs / No. Genes, the rate of ARGs in all genes. The number of ARGs in public database (e.g., UniProt) is obtained by text mining. All genes as shown in this table are based on length filtration of the original 240,089,111 genes obtained from 815 microbiome samples, as provided in **Table S5**.
