## Supplementary material for "Ontology-aware deep learning for antibiotic resistance gene prediction: novel function discovery and comprehensive profiling from metagenomic data": Supplementary Table S11.docx

**Supplementary Table 11. The binding affinity of protein–ligand complexes using the top five pockets.**

| **ID** | **Pocket1** | **Pocket2** | **Pocket3** | **Pocket4** | **Pocket5** |
| --- | --- | --- | --- | --- | --- |
| Candi_60363_1 | -7.6 | -7.6 | -7.5 | **-7.7** | -7.5 |
| AHE40557.1 | -8.1 | **-8.4** | -7.3 | -8.1 | -8.3 |
| Negative control | -6.0 | -5.9 | -5.8 | **-6.1** | -6.0 |

Note: The values are binding affinity (kcal/mol).
