## Supplementary figures and images for "Ontology-aware deep learning for antibiotic resistance gene prediction: novel function discovery and comprehensive profiling from metagenomic data"

### Supplementary Figure S1.pdf

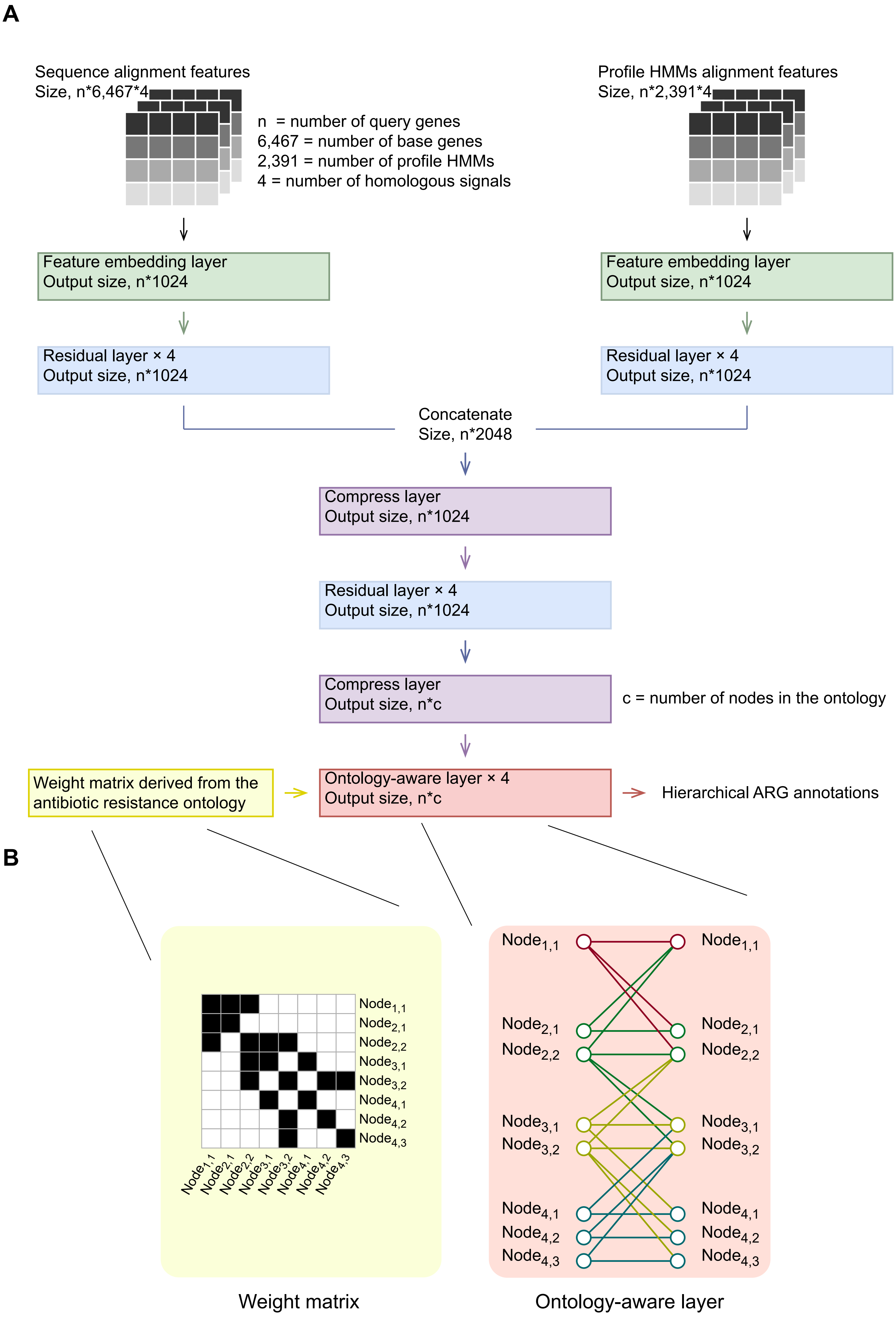

### Supplementary Figure S2.pdf

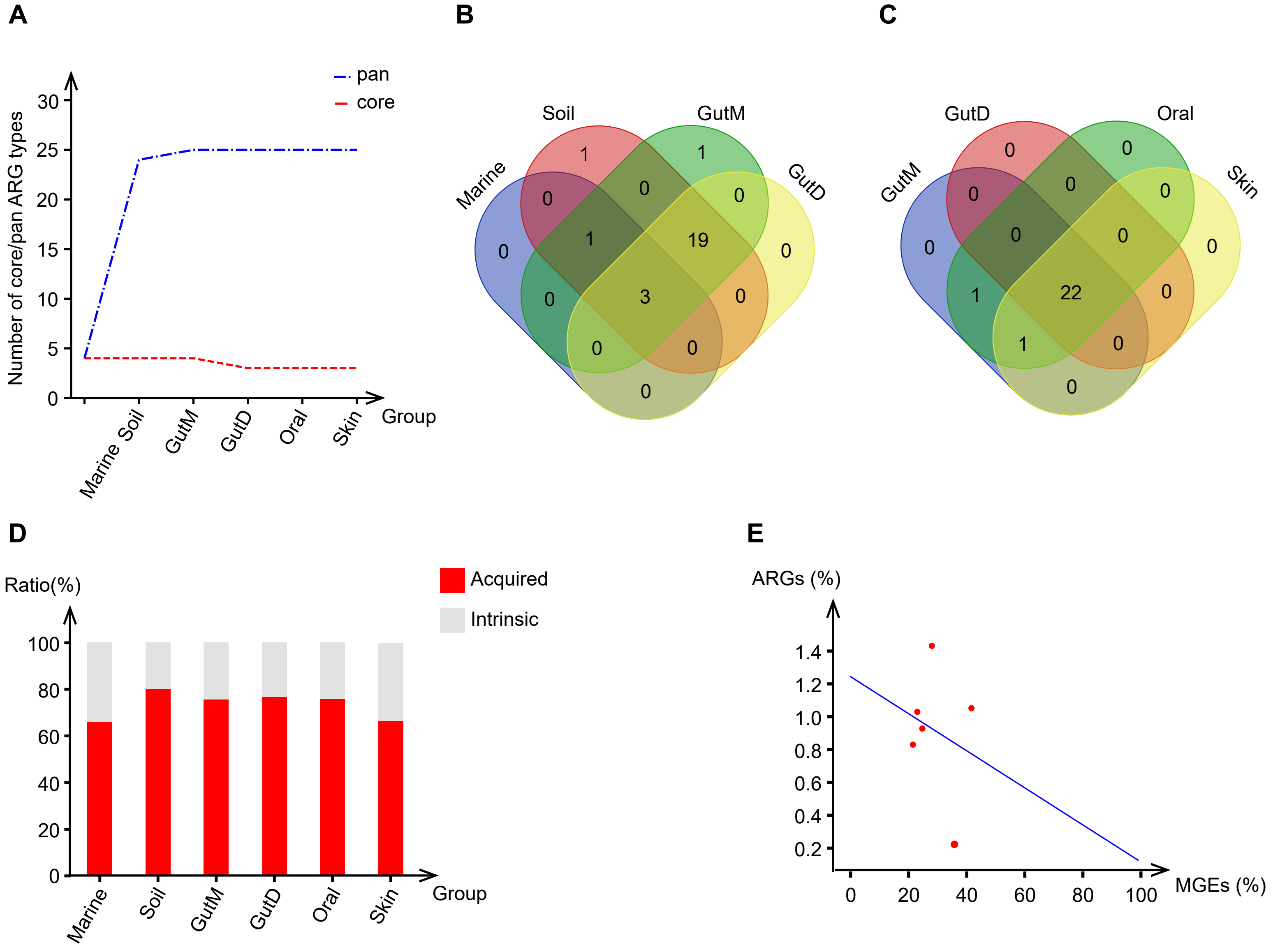

### Supplementary Figure S3.pdf

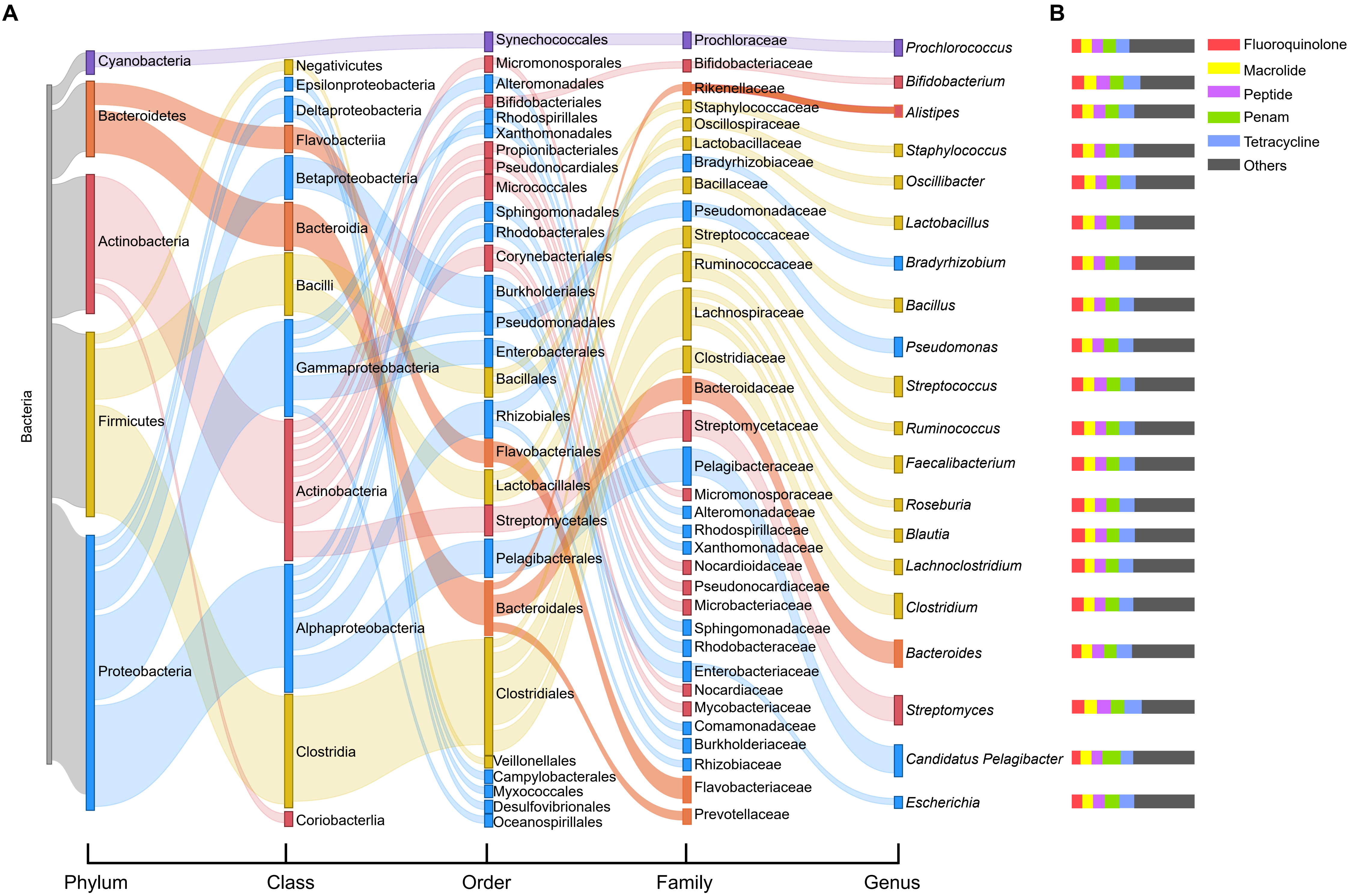

### Supplementary Figure S4.pdf

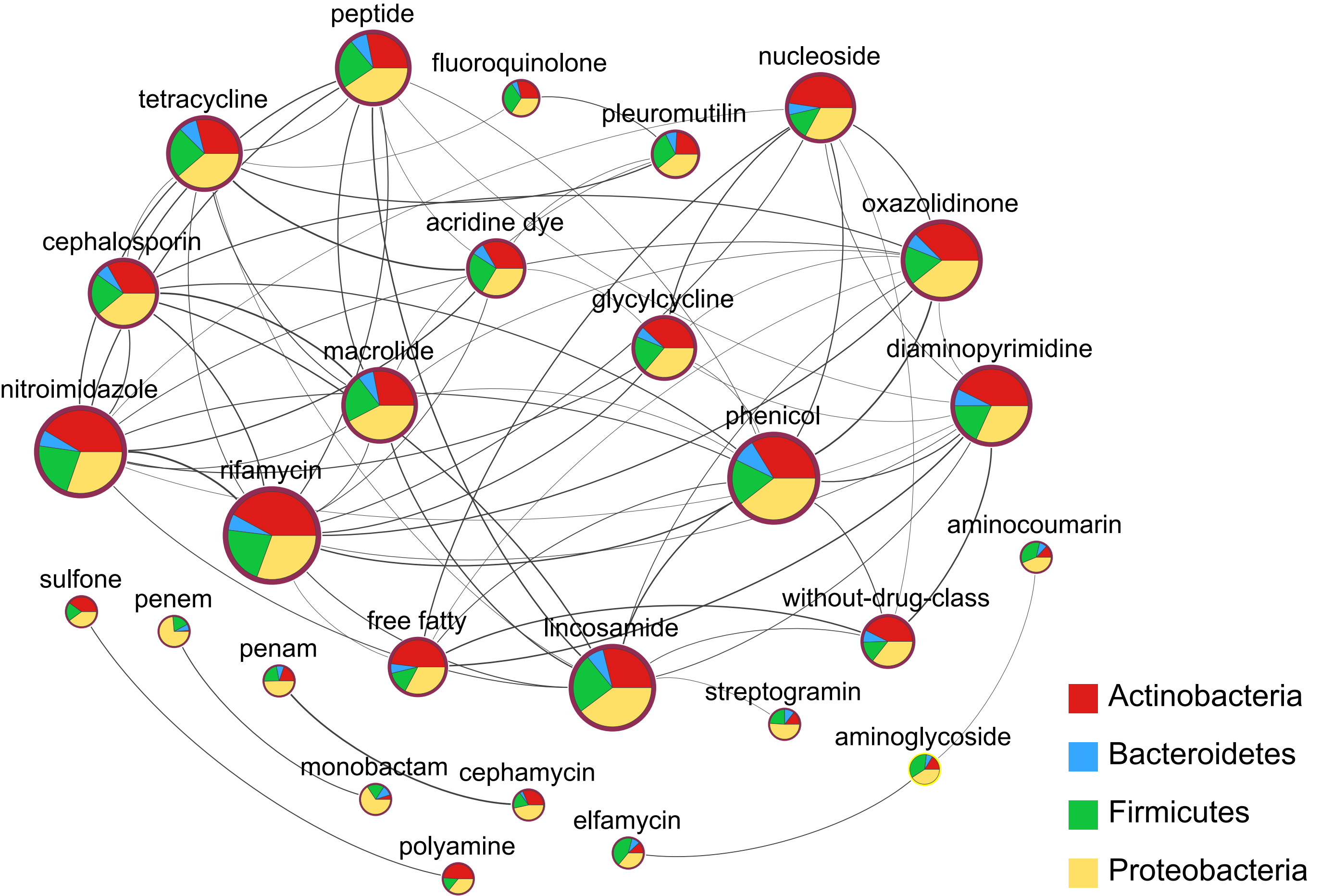
